# Evolutionary insights into bilin biosynthesis: Functional characterization of pre-PcyA enzymes

**DOI:** 10.64898/2026.05.08.723791

**Authors:** Federica Frascogna, Nathan C. Rockwell, Gunhild Layer, Nicole Frankenberg-Dinkel

## Abstract

Biosynthesis of the linear tetrapyrrole phycocyanobilin (PCB) by the ferredoxin-dependent bilin reductase (FDBRs) PcyA is essential for light-harvesting and regulatory processes in diverse photosynthetic organisms, yet its evolutionary origins are not fully understood. PcyA evolved from pre-PcyA proteins found in diverse bacteria. Three lineages of pre-PcyA proteins were identified: Pre-1, Pre-2 and Pre-3. Using an *in vivo* co-expression assay, Pre-2 and Pre-3 proteins were shown to be active FDBRs that did not synthesize PCB, whereas Pre-1 activity was apparently low. In refining these results, we noted a discrepancy between phycoerythrobilin populations generated by Pre-3 and by the distantly related FDBR PebS. We therefore examined the properties of pre-PcyA enzymes *in vitro*, using an updated pre-PcyA phylogeny to select an alternative pre-1 target. Biochemical analyses revealed that Pre-1 and Pre-2 catalyze the two-electron reduction of biliverdin (BV) to *3E*-phytochromobilin (*3E*-PФB), in contrast to the known synthesis of *3Z*-phytobilins by other FDBRs. Pre-3 can also carry out an additional two-electron reduction to yield *3E*-phycoerythrobilin (*3E*-PEB), again distinct from the *3Z*-PEB produced by PebS. We then used comparative sequence and structure analysis to target candidate catalytic residues for site-directed mutagenesis. Variant Pre-1 exhibited altered product stereochemistry, but no effects on Pre-2 were observed and Pre-3 variants unexpectedly gained the ability to bind cyclic tetrapyrroles. These findings underscore the plasticity and promiscuity of this enzyme family. Together, this work illustrates how the flexible catalytic potential of ancestral enzymes shaped the evolution and diversification of bilin biosynthetic pathways.

## Introduction

Bilins are open chain tetrapyrrole molecules, widespread in nature and often regarded as “pigments of life”, alongside other tetrapyrroles (Battersby, 1985). These pigments serve diverse functions, including involvement in light harvesting and chlorophyll synthesis in oxyphototrophs as well as light sensing in a broad range of organisms (Rockwell & Lagarias, 2020; Zhang et al., 2021; Richardson & Campbell, 2022; Bryant & Gisriel, 2024; Zhao et al., 2025). Bilin biosynthesis is both ubiquitous and essential in oxygenic photosynthetic organisms (Alvey et al., 2011; Duanmu et al., 2013). Consistent with this, all oxyphototrophs retain the heme oxygenase (HO) and ferredoxin-dependent bilin reductase (FDBR) enzymes responsible for bilin biosynthesis, regardless of the presence of bilin-based light harvesting or photoreceptors (Rockwell & Lagarias, 2017; Rockwell et al., 2023). However, different clades of photosynthetic organisms have different FDBRs and hence may synthesize different bilins from the common biliverdin IXα (BV) precursor produced from heme by HO.

FDBRs group into three major lineages: PcyA/PcyX, PebA/PebS/PUBS, and PebB/HY2 (Rockwell & Lagarias, 2017; Ledermann et al., 2018;)Frascogna et al., 2026;. The PcyA/PcyX lineage includes sequences from cyanobacteria, from prasinophyte and glaucophyte algae, and from some phages. PcyA enzymes catalyze a 4-electron reduction of BV to yield phycocyanobilin (PCB), which is essential for the normal function of the light-harvesting phycobilisome of cyanobacteria, red algae, and glaucophytes. PcyX is confined to phages and instead catalyzes the 4-electron reduction of BV to phycoerythrobilin (PEB). The PebA/PebS/PUBS and PebB/HY2 groups are present in cyanobacteria, rhodophytes, streptophytes, and a broad range of eukaryotic algae with plastids derived from red algae, while the PebA/PebS/PUBS group is also found in cyanophages (Rockwell & Lagarias, 2017; Baeuerle et al., 2026). PebA and PebB synthesize PEB from BV in two steps: PebA first carries out a 2-electron reduction at the 15,16-double bond of BV to yield 15,16-dihydrobiliverdin (15,16-DHBV), and PebB then carries out another 2-electron reduction at the bilin A-ring to form PEB. PebS is confined to cyanophages and synthesizes PEB via the same 4-electron reduction performed by PcyX. PUBS also carries out a 4-electron reduction of BV. HY2 is found in streptophytes, and its behavior is somewhat more complex. In streptophyte algae, it carries out the same 4-electron reduction of BV as PcyA, producing PCB. In land plants, HY2 again uses BV as a substrate but carries out the A-ring reduction performed by PebB to synthesize phytochromobilin (PΦB).

Bilin-based phycobilisome antenna complexes are used by most cyanobacteria for light harvesting, a process that relies on PCB to relay energy to chlorophylls. Phycobilisomes are also found in the Gloeobacterales, early-branching cyanobacteria that lack thylakoid membranes, and can provide more complex cyanobacteria with adaptable light-harvesting apparatus that can be optimized for the ambient light environment (Gan et al., 2014; Hirose et al., 2019; Soulier et al., 2020). One might thus expect that the earliest bilin in cyanobacteria would be PCB, and hence the earliest cyanobacterial FDBR would be PcyA. Consistent with this, the *pcyA* gene is essential in the model cyanobacterium *Picosynechococcus sp*. PCC 7002 (ex *Synechococcus*; (Alvey et al., 2011; Komárek et al., 2020). In contrast, *pebA* and *pebB* are restricted to PEB-producing cyanobacteria (Frankenberg et al., 2001). However, the ongoing sequencing of new genomes and metagenomes implicates a more complex evolutionary history of this enzyme family. Notably, almost a decade ago, *pcyA* was identified in the photosynthetic alphaproteobacterium *Bradyrhizobium* sp. ORS278, and phylogenetic analyses revealed that these PcyA sequences are not derived from cyanobacterial ancestors (Jaubert et al., 2007; Ledermann et al., 2018). These proteins, along with the more recently identified PcyX, form an FDBR lineage known as the AX clade (Ledermann et al., 2016; Rockwell et al., 2023). More recently, FDBR-like genes were discovered in heterotrophic bacteria. These sequences, referred to as pre-PcyAs, clustered into three distinct lineages: Pre-1, Pre-2, and Pre-3 (Rockwell et al., 2023). Pre-1 and Pre-2 sequences are found in both metagenome-assembled genomes (MAGs) and cultured organisms, whereas Pre-3 is exclusively found in MAGs to date. The distribution of these clades also varies: Pre-1 is widespread across multiple bacterial phyla, including beta- and gamma-proteobacteria, Acidobacteria, Planctomycetes, and Verrucomicrobia, whereas Pre-2 and Pre-3 are primarily restricted to Myxococcota. Pre-1 and Pre-3 genes are typically organized in clusters alongside open reading frames encoding a heme oxygenase (ho) and bilin-biosynthesis-associated globins (BBAGs), whereas Pre-2 is typically associated with bilin- and B12-binding proteins homologous to the two chromophore-binding domains found in photocobilin photoreceptors (Rockwell et al., 2023; Zhang et al., 2024). Pre-PcyA proteins show considerable divergence from authentic PcyA proteins, especially in their active sites. The 4-electron reduction of BV catalyzed by PcyA begins with the reduction of the C18^1^-C18^2^ exo-vinyl group to yield the transient intermediate 18^1^,18^2^-dihydrobiliverdin (18^1^,18^2^-DHBV), followed by the reduction of the A-ring diene system (Frankenberg & Lagarias, 2003; Tu et al., 2004; Tu et al., 2007; Hagiwara et al., 2010). This activity strictly depends on a catalytic triad consisting of Glu^76^, His^88^, and Asp^105^. The reaction is initiated by Asp^105^-mediated protonation of BV, with Glu^76^ facilitating a second protonation at the D-ring exo-vinyl group. Finally, His^88^ mediates D-ring reduction and subsequent proton transfer via a relay system from the bulk solvent (Tu et al., 2007). In contrast, the pre-PcyA lineages exhibit systematic variations in these critical residues. Notably, equivalents to Glu76 and His88 are entirely absent in pre-PcyA proteins. Asp105 is typically equivalent to Glu in Pre-1 but is present in both Pre-2 and Pre-3. Consistent with this variation, characterization of these proteins using *in vivo* co-expression assays did not demonstrate synthesis of PCB by representatives of each clade (Rockwell et al., 2023). Instead, Pre-1 and Pre-2 were found to catalyze a 2-electron reduction of BV A-ring to PФB, albeit with low activity in the chosen Pre-1 protein (POZ53557 from *Methylovulum psychrotolerans*). The pre-3 protein MBL9008304 carried out this reaction as well, but it also carried out subsequent reduction of the 15,16-double bond to yield a mixture of PФB and PEB. We here provide the first *in vitro* characterization of pre-PcyA enzymes. Further characterization of BBAG preparations incorporating different bilins in co-expression assays revealed discrepancies between PEB synthesized by Pre-3 and by PebS, in contrast to consistent results obtained for PФB synthesized by Pre-2 and Pre-3. We therefore purified both of these FDBRs, along with an alternative Pre-1 enzyme chosen using an updated phylogenetic analysis. All three pre-PcyA proteins generated *3E*-PФB, with Pre-3 also producing *3E*-PEB. This contrasts with synthesis of *3Z* phytobilins by other FDBRs. Attempts to modulate the catalytic activity of pre-FDBRs by site-directed mutagenesis produced variable results: Pre-1 was engineered to produce *3Z*-PФB, but Pre-2 catalysis was unchanged and Pre-3 acquired the capacity to bind cyclic tetrapyrroles (porphyrins). Together, these findings highlight the diversity of pre-PcyAs and the broader enzymatic plasticity of the FDBR superfamily.

## Results

### Distinct PEB populations formed by PebS and by the pre-3 MBL9008304

In previous work, the pre-3 protein MBL9008304 was characterized using *in vivo* co-expression assays with two acceptor proteins: the model cyanobacterial phytochrome Cph1 and its own cognate BBAG, MBL9008307 (Rockwell et al., 2023). This work resulted in preparations of Cph1 and MBL9008307 containing mixtures of PФB and PEB. The behavior of Cph1 with both PФB and PEB chromophores as individual chromophores is well understood (Murphy & Lagarias, 1997; Yeh et al., 1997; Borucki et al., 2003; Fischer et al., 2005). To better characterize PФB and PEB populations of MBL9008307, we purified this BBAG after recombinant co-expression either with the previously characterized pre-2 protein CAP_1520 from *Chondromyces apiculatus* (producing PФB; BBAG+CAP) or with PebS (producing PEB via 15,16-DHBV; 8307BBAG+PebS). BBAG+CAP (Pre-2) exhibited bilin absorption peaks at 372 nm and 648 nm, along with a smaller protein absorption peak at 280 nm (Fig. 1A and Table 1). The fluorescence excitation spectrum of BBAG+CAP exhibited a maximum at 647 nm, with an emission peak at 663 nm (Fig. 1B). These values were very similar to those obtained after co-expression with the Pre-3 MBL9008304 (BBAG+8304; Fig. 1C and Table 1). The absorption spectrum of BBAG+PebS exhibited a major peak at 552 nm, with additional peaks at 386 nm and 302 nm (Fig. 1D). The latter peak was partially overlapping the protein absorption peak, which was consequently red-shifted to 284 nm. Similar peaks at 300-310 nm have previously been reported for PEB adducts of phytochromes (Murphy & Lagarias, 1997). A small shoulder at approximately 595 nm was also present, which could arise from a small amount of 15,16-DHBV. Fluorescence excitation of this BBAG+PebS preparation peaked at 551 nm, with an emission maximum at 566 nm (Fig. 1E). These fluorescence wavelengths were blue-shifted relative to those seen for BBAG+8304 (Fig. 1C and Table 1), as was the PEB absorption peak (Fig. 1F). These results thus demonstrated that the PФB populations of BBAG MBL9008307 as produced by Pre-2 or Pre-3 enzymes were equivalent, but that the population assigned to PEB after co-expression with Pre-3 was spectroscopically distinct from that seen after co-expression with PebS. It thus seemed possible that Pre-3 was either producing a different PEB isomer or was not producing PEB at all. We therefore sought to directly characterize the reactions of pre-PcyA proteins with BV *in vitro*.

**Figure 1.**
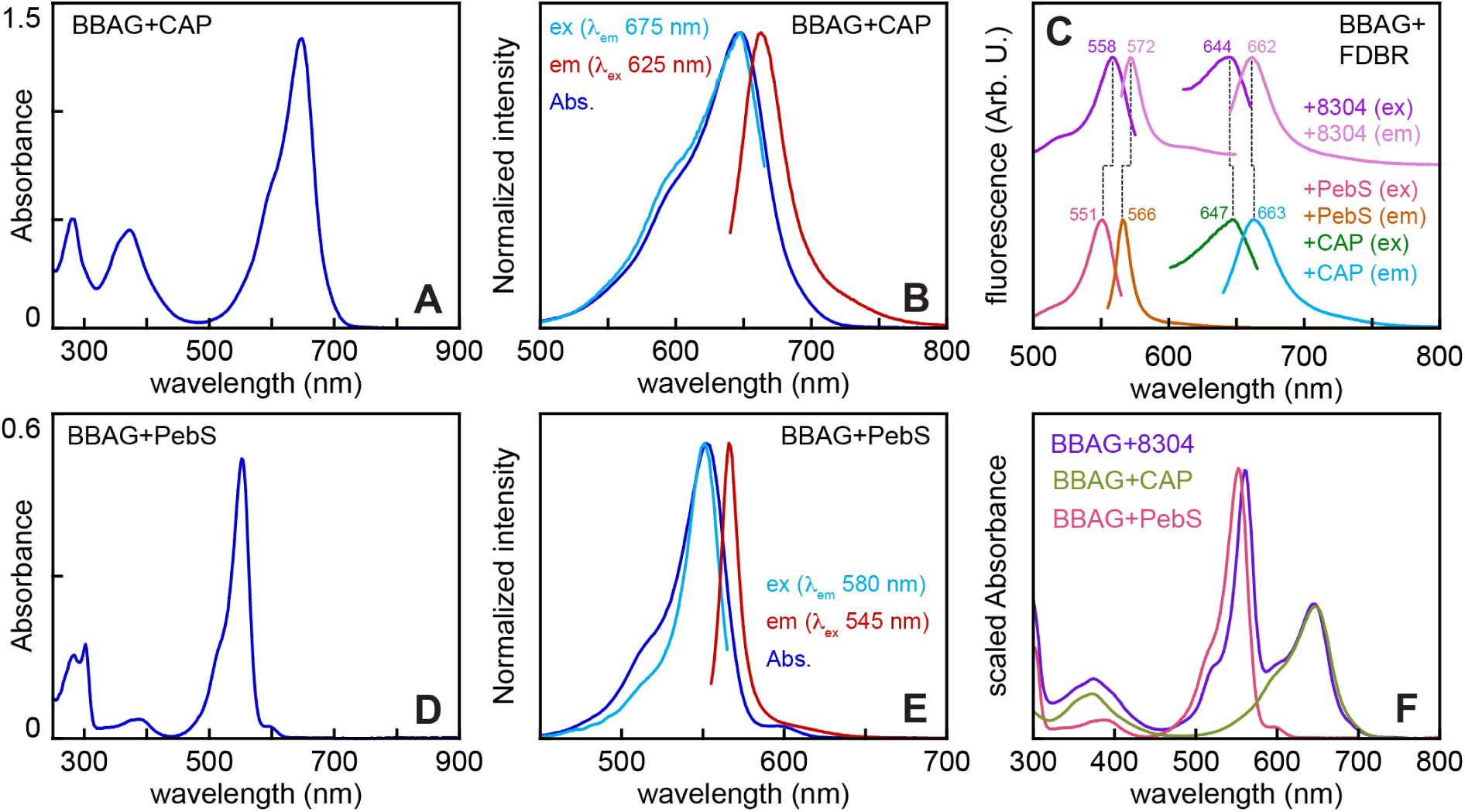
Characterization of BBAG MBL9008307 after co-expression with different FDBRs. (A) The absorption spectrum is shown for BBAG MBL9008307 after co-expression with pre-2 CAP_1520 (BBAG+CAP), which produces PФB. (B) Normalized absorption (dark blue), fluorescence excitation (cyan), and fluorescence emission (red) spectra are shown for BBAG+CAP. (C) Fluorescence excitation and emission spectra are shown for BBAG MBL9008307 with the indicated FDBRs. The pre-3 FDBR MBL9008304 is the cognate FDBR for this BBAG, producing both PФB and PEB. Peak wavelengths are indicated, and dashed lines indicate comparable spectra. (D) The absorption spectrum is shown after co-expression with PebS (BBAG+PebS), which produces PEB via 15,16-DHBV. (E) Normalized absorption (dark blue), fluorescence excitation (cyan), and fluorescence emission (red) spectra are shown for BBAG+PebS. (F) Scaled absorption spectra are shown for BBAG+CAP (olive), BBAG+PebS (coral), and BBAG+8304 (purple; after co-expression with MBL9008304) (Rockwell et al., 2023).

**Table 1:**
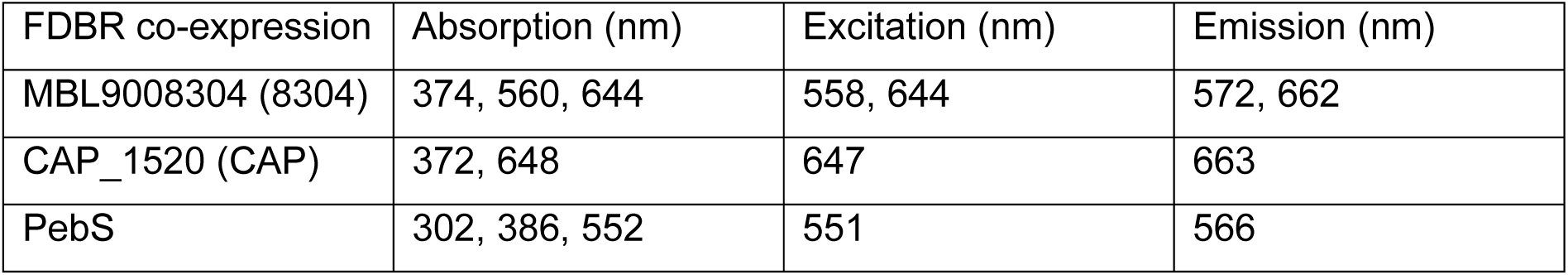
Peak wavelengths for native BBAG MBL9008307.

### Selection of enzymes for purification and characterization

Previous examination of FDBRs demonstrated robust activity for the pre-2 enzyme CAP_1520 and the pre-3 enzyme MBL9008304 (Rockwell et al., 2023). However, the pre-1 candidate POZ53557 from *Methylovulum psychrotolerans* exhibited little activity, with apparent formation of only trace amounts of PФB in the presence of the model phytochrome Cph1 as an acceptor protein (Rockwell et al., 2023). This contrasts with the robust activity of its cognate HO in similar assays (Rockwell et al., 2022; Rockwell et al., 2023). We therefore sought an alternative sequence from the pre-1 clade that might offer robust catalytic activity. More recent phylogenetic analysis of FDBRs (Frascogna, Rockwell et al., 2026) identified additional candidate lineages, so we updated that analysis (Fig. S1) to identify candidate proteins meeting three criteria: first, reliable assignment to the pre-1 lineage; second, a representative chromosomal context including HO, BBAG, V4R, and FeS proteins (Rockwell et al., 2023); and, third, representative active-site residues for the pre-1 lineage. This analysis largely matches recent work (Frascogna, Rockwell et al., 2026); major differences are in the placement of uncharacterized lineages that were also not robustly placed in the previous work. Within the pre-1 clade (Fig. 2A), the metagenomic sequence MCC5789364 satisfied these three criteria. It has been placed within this clade in all recent analyses and is closely related to POZ53557, supporting a robust assignment as a pre-1 enzyme. MCC5789364 is encoded on a scaffold also encoding candidate BBAG, FeS, and V4R proteins (Fig. 2B). However, there is no HO on this scaffold (JAFIGF010000120 in NCBI assembly GCA_020831535.1). BLAST (Altschul et al., 1997) searches identified two candidate HO sequences on different scaffolds within this metagenome assembled genome (MAG). One of the two, MCC5789513, is near the edge of its scaffold (JAFIGF010000136). Both scaffolds end in the middle of open reading frames, so it seemed plausible that this HO might be the cognate HO but was separated from the rest of the cluster during the assembly process due to a scaffold break. In support of this hypothesis, the two fragmentary proteins could be matched to a single protein in a different assembly from the same experimental series (Ataeian et al., 2022). This protein (MCH8539434) is encoded on scaffold JAIUMF010000008 in assembly GCA_022565675.1, with both MAGs assigned as *Opitutales* spp. This alternative scaffold contains identical matches to both the candidate FDBR MCC5789364 and the candidate HO MCC5789513 (MCH8539424 and MCH8539435, respectively; Fig. 2B). Taken together, it seems likely that MCC5789364 is a pre-1 FDBR found as part of a complete cluster of associated proteins. Based on the sequence alignment underlying our phylogenetic analysis, MCC5789364 also exhibits candidate active-site residues typical of a pre-1 enzyme (Fig. 2A). For example, the critical Asp105 in PcyA (Tu et al., 2007; Joutsuka et al., 2023) instead corresponds to Glu in these proteins, including both MCC5789364 and POZ53557 (Fig. 2C and Table 2). His88 in PcyA most frequently corresponds to Thr or Asn in pre-1. POZ53557 has Thr at this position, whereas MCC5789364 has Asn (Fig. 2C). The third residue known to play a key role in PcyA catalysis, Glu76, does not correspond to a conserved consensus residue in pre-1 proteins (Hagiwara et al., 2010) (Fig. 2A). Asp256 in Arabidopsis HY2 is important for A-ring reduction (Tu et al., 2008; Sugishima et al., 2020; Frascogna et al., 2023), the same reaction proposed to occur in pre-1 and known to occur in other pre-PcyA proteins (Rockwell et al., 2023). This residue typically corresponds to Ser in pre-1 proteins, including MCC5789364 but not POZ53557 (Fig. 2C). MCC5789364 thus exhibits the typical features of a pre-1 FDBR and seemed to provide a suitable candidate for *in vitro* characterization of Pre-1, with CAP_1520 and MBL9008304 providing corresponding candidates for Pre-2 and Pre-3.

**Figure 2.**
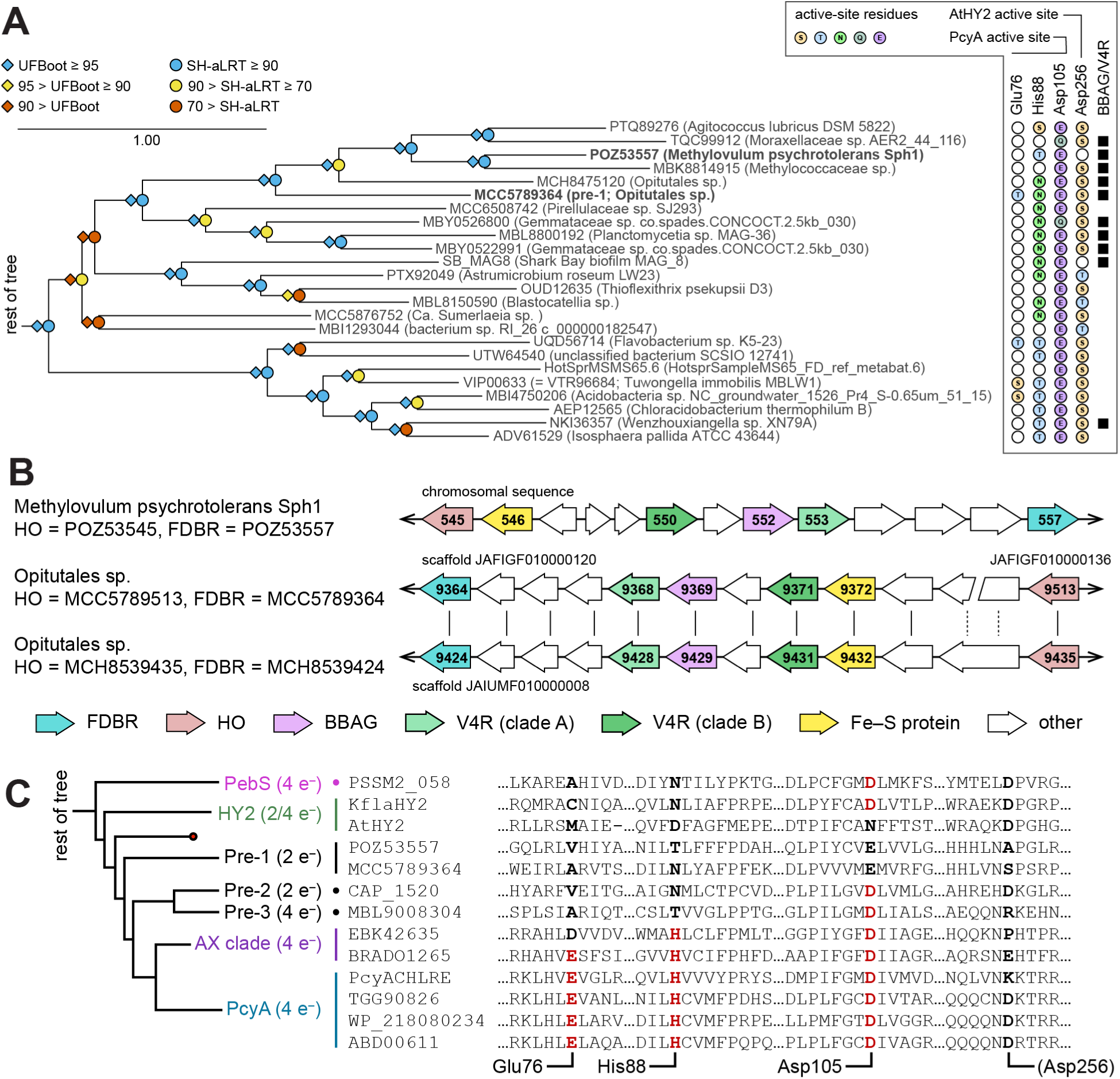
Identification of an alternative pre-1 enzyme for characterization. (A) The local region around the previously examined pre-1 POZ53557 is shown, with the full phylogenetic tree presented in Fig. S1. Residues corresponding to known catalytic residues in *Synechocystis* PcyA or *Arabidopsis* HY2 are indicated; cases where no apparent enrichment was found are left as open white circles, and other residues are indicated. Proteins found in the context of clusters with BBAGs and V4R proteins are indicated with black squares. Experimentally characterized POZ53557 (Rockwell et al., 2023) and MCC5789364 (this work) are indicated in bold face. (B) The local chromosomal context is shown for POZ53557, MCC5789364, and MCH8539424, with metagenomic scaffolds indicated. The latter two FDBRs are identical in protein sequence. The MCC5789364 cluster is apparently broken by a scaffold break, resulting in the protein sequence equivalent to MCH8539434 appearing at the edges of two scaffolds (dashed vertical lines). (C) Candidate active-site residues (bold face) are shown for selected FDBRs. Conservation of Glu76, His88, and Asp105, essential for synthesis of PCB by PcyA, is indicated in red. Phylogenetic placement within the tree of Fig. S1 is indicated on the left, along with the presence of 2-electron or 4-electron reductases within each clade.

**Table 2:** Amino acid equivalents in selected FDBRs. Residues included in the active site analysis (Figs. 2A & C; Fig. 6) are indicated relative to PcyA from *Synechocystis* sp. PCC 6803 (Glu76, His88, and Asp105; NCBI accession slr0116 and PDB accession 2D1E) or relative to HY2 from *Arabidopsis thaliana* (Asp256; NCBI accession OAP03522). Reference residues are in **bold**, and residues targeted for site-directed mutagenesis are in ***bold italic***. The NCBI accession for KflaHY2 is kfl00177_0010, and numbering is based on the full-length protein (including plastid targeting signals). Asp208 and As328 correspond to Asp122 and Asp242 in the previously described expression construct (Frascogna et al., 2023).

| FDBR type | Protein | Glu76 | His88 | Asp105 | (Asp256) |
| --- | --- | --- | --- | --- | --- |
| PcyA | *SynPcyA* | **Glu76** | **His88** | **Asp105** | Asp220 |
| PcyA | *NosPcyA* | Glu73 | His85 | Asp102 | Asp217 |
| PcyA | ABD00611 | Glu71 | His83 | Asp100 | Asp216 |
| PcyA | BRADO1265 | Glu71 | His83 | Asp100 | Glu211 |
| PcyX | EBK42635 | Asp72 | His86 | Asp103 | Pro216 |
| Pre-1 | MCC5789364 | Ala63 | Asn74 | **Glu91** | Ser202 |
| Pre-2 | CAP_1520 | Val63 | Asn74 | **Asp91** | Asp207 |
| Pre-3 | MBL9008304 | Ala66 | Thr79 | **Asp96** | Arg215 |
| HY2 | KflaHY2 | Cys179 | Asn191 | Asp208 | Asp328 |
| HY2 | AtHY2 | Met115 | Asp116 | Asn133 | **Asp256** |
| PebS | PSSM2_058 | Ala77 | Asn88 | Asp105 | Asp206 |

### Biochemical investigation of pre-PcyA activities

To trace the origin of FDBRs, we recombinantly produced and purified these three representative pre-PcyA enzymes and examined their activity. The activity of Pre-1 was investigated employing equimolar amounts of enzyme and BV, as the potential substrate (Fig. 3A). The incubation of the putative reductase with BV resulted in an intensely teal-colored complex with an absorbance maximum at ∼ 650 nm and a small shoulder at ∼ 600 nm, suggesting the ability of the protein to bind the substrate (Fig. 3A, blue curve). Additionally, the presence of a shoulder at ∼ 730 nm suggested the presence of protonated BV (Tu et al., 2004; Tu et al., 2007; Tu et al., 2008; Busch et al., 2011). Upon initiation of the reaction by addition of the NADPH-regenerating system (Fig. 3A, dashed curve), no immediate effect could be observed. However, as the assay progressed, the main peak gradually shifted toward ∼ 670 nm and, concomitantly, an increase in absorbance at ∼ 730 nm was detected, implying the formation of substrate radical intermediates (Fig. 3A, red curve). The reaction product was isolated from the reaction mix and identified via HPLC (Fig. 3B). The analysis revealed the formation of a main product with a retention time of 15.5 min (Fig. 3B, Pre-1), not overlapping with any available bilin standard (Fig. 3B). To elucidate its identity, further investigation was conducted using coupled phytochrome assembly assays employing apo-Cph1 from *Synechocystis* sp. PCC 6803 (apo-*Syn*Cph1: Fig. 3C), which incorporates PCB as its natural chromophore (Yeh et al., 1997), and apo-BphP from *Pseudomonas aeruginosa* (apo-*Pa*BphP), which incorporates BV (Tasler et al., 2005). When the unidentified product was incubated with apo-*Pa*BphP, no characteristic difference spectrum was obtained (data not shown), consistent with the absence of a photoactive adduct and hence with the absence of the A-ring endo-vinyl group. In contrast, the incubation with apo-*Syn*Cph1 resulted in a difference spectrum nearly identical to the one obtained with PФB (Kohchi et al., 2001; Sawers et al., 2004; Muramoto et al., 2005; Mukougawa et al., 2006; Zhao et al., 2007; Lu et al., 2017), suggesting that the produced bilin might be 3*(E)*-PФB. The PФB used as standard for the HPLC analysis, was obtained from an anaerobic bilin-reductase activity assay performed using the HY2 from *Arabidopsis thaliana*. Previously characterized FDBRs such as *At*HY2 and PebS predominantly generate the 3*(Z)* bilin isomer (Frankenberg et al., 2001; Dammeyer et al., 2008). Hence, production of 3*(E)*-PФB by Pre-1 provides the first reported instance of an FDBR exhibiting a preference for the 3*(E)* isomer *in vitro* under anaerobic conditions.

**Figure 3.**
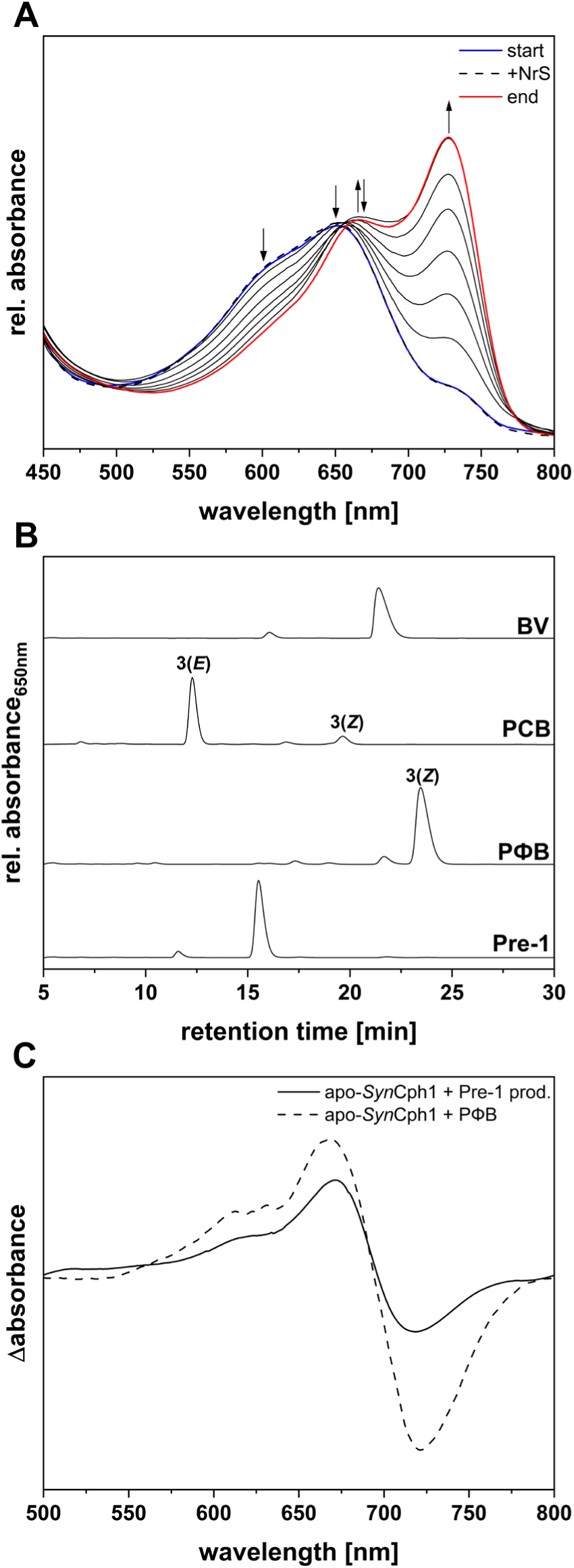
Investigation of the activity of recombinant Pre-1 and identification of reaction products. (A) Time-dependent spectra of an anaerobic bilin reductase activity assay using recombinant Pre-1 and BV as the substrate. The total reaction time was 20 minutes and spectra were recorded at 30 seconds intervals, with representative timepoints shown for clarity. Arrows indicate the absorbance changes during the reaction. The blue spectrum corresponds to the “binding spectrum”, recorded upon incubation of BV with Pre-1. The dashed spectrum is the first recorded after initiating the reaction with the NADPH-regenerating system (NrS), and the final spectrum is red. A 50-point Savitzky-Golay filter was applied to smooth the curves. (B) HPLC analysis of the reaction products (Pre-1). The products were separated on a reverse-phase 5 μm C18 Luna column (Phenomenex), with a mobile phase consisting of 50% acetone (v/v) and 50% 20 mM aqueous formic acid (v/v), flowing at 0.6 mL/min. Absorbance was continuously monitored at 650 nm. BV, biliverdin IXα standard; PCB, phycocyanobilin standard; PΦB, phytochromobilin standard obtained from an *Arabidopsis thaliana* HY2 assay with BV. (C) Difference spectra of coupled phytochrome assembly assays employing apo-*Syn*Cph1 incubated with a PФB standard or the product of the Pre-1 reaction (Pre-1 prod.). Spectra were recorded after incubation for 3 min with red light (636 nm – Pfr spectrum) and following incubation for 3 min with far-red light (730 nm – Pr spectrum). Difference spectra were calculated by the subtraction of the Pfr from the Pr spectrum. The apo-*Syn*Cph1 + PФB difference spectrum (dashed line) was smoothed applying a 10 pt. Savitzy-Golay filter. The apo-*Syn*Cph1 + Pre-1 prod. difference spectrum (solid line) was smoothed applying a 40 pt. Savitzky-Golay filter and the signal was 3x enhanced.

The activity of Pre-2 (CAP_1520) was investigated using the same approach (Fig. 4A), assuming that BV was its natural substrate based on its robust synthesis of PФB in co-expression assays with Cph1 (Rockwell et al., 2023) and MBL9008307 (Fig. 1). Incubation of Pre-2 with BV resulted in the formation of a complex with an absorbance maximum at ∼ 670 nm (Fig. 4A, blue curve). The start of the reaction had little effect on the initial absorbance peak (Fig. 4A, dashed curve). The reaction proceeded at a slow turnover rate, ultimately yielding a product absorbing at 645 nm without accumulation of the far-red species seen with Pre-1 (Fig. 4A, red curve). HPLC analysis of the isolated product indicated that the reaction was incomplete, as residual BV was detected at a retention time of ∼ 21 min, while the main product eluted at 14.5 min (Fig. 4B, Pre-2). This reaction product matched the product of the Pre-1 reaction (Fig. 4B, Pre-1), with slight variation in retention times due to slight differences in the mobile phase. This implicates production of 3*(E)*-PФB by both Pre-1 and Pre-2, which we again supported using phytochrome assembly assays with apo-*Syn*Cph1 and apo-*Pa*BphP. Once again, only the incubation of the product with apo-*Syn*Cph1 led to the formation of a photoactive adduct, again exhibiting the characteristic PФB-Cph1 photocycle (Fig. 4C). We therefore conclude that both Pre-1 and Pre-2 catalyze the conversion of BV to 3(*E*)-PФB. Like Pre-1, Pre-3 (MBL9008304) is encoded in a gene cluster alongside a heme oxygenase; it also exhibited robust activity in co-expression assays, like Pre-2 (Rockwell et al., 2023). We therefore used BV as the substrate in an anaerobic bilin-reductase activity assay. An initial assay revealed a color shift of the reaction mix from teal to antique pink, consistent with the expected conversion of BV to PEB. This prompted a time course assay, in which samples were withdrawn from the mix after 5, 9 and 20 minutes from the start of the reaction to gain more insights into the reaction course (Fig. 5A). The incubation of Pre-3 with the substrate led to the formation of a complex with dual absorbance peaks at ∼ 660 nm and ∼ 710 nm, with the latter likely corresponding to BVH^+^ (Fig. 5A, blue curve). Upon initiation of the reaction (Fig. 5A, dashed curve), no noticeable changes were observed, and the initial absorbance peaks remained stable for the first 5 minutes (Fig. 5A, cyan curve). By 9 min, they transitioned into a single peak at ∼ 690 nm (Fig. 5A, pink curve), whereas the final 20 min product exhibited an additional peak at ∼ 560 nm (Fig. 5A, red curve). HPLC analysis of these three products revealed an initial conversion of BV to 3(*Z*)-PФB as monitored at 650 nm (Fig. 5B, 5 min). This peak decayed at later times. At 20 minutes, we clearly detected 3(*E*)-PEB (Fig. 5C). The conversion of 3(Z)-PФB to 3(*E*)-PEB would require formation of either 3(*Z*)-PEB or 3(*E*)-PФB as an intermediate. In monitoring the reaction at 650 nm, we observed increasing signal at 17 min retention time during the course of the reaction (Fig. 4B), so we favor formation of 3(*E*)-PФB as an intermediate followed by subsequent reduction of the 15,16-double bond to yield 3(*E*)-PEB. These results thus confirmed the reaction pathway inferred from co-expression assays: Pre-3 carries out the first reported example of a 4-electron reduction of BV to PEB with PФB serving as the intermediate rather than 15,16-DHBV.

**Figure 4.**
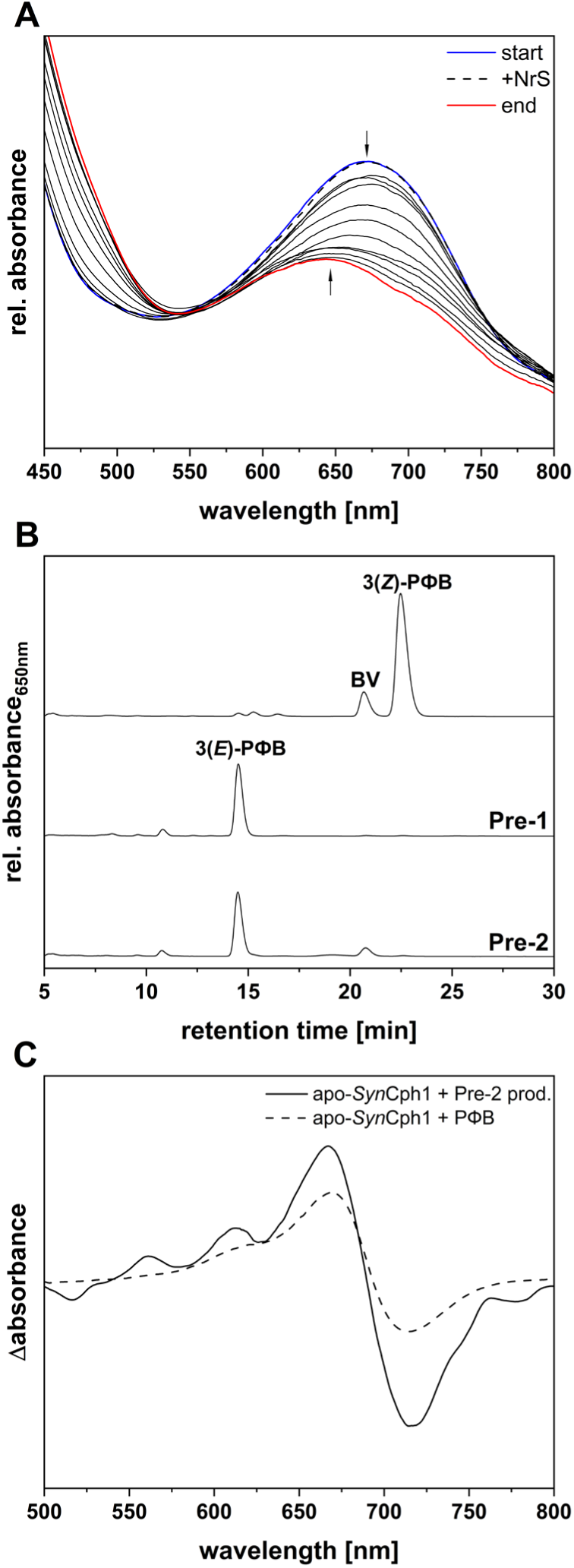
Investigation of the activity of recombinant Pre-2 and identification of reaction products. (A) Time-dependent spectra of an anaerobic bilin reductase activity assay using recombinant Pre-2 and BV as the substrate. The total reaction time was 30 minutes and spectra were recorded at 30 seconds intervals, although, for clarity, only informative spectra are shown. The arrows follow the absorbance changes during the reaction. The blue spectrum corresponds to the “binding spectrum”, recorded upon incubation of BV with Pre-2. The dashed spectrum represents the first recorded after initiating the reaction with the NrS. The black spectra indicate the most relevant ones recorded during the assay, while the final spectrum is red. An 80-points Savitzky-Golay filter was applied to smooth the curves. (B) HPLC analysis of the reaction products (Pre-2). The products were separated on a reversed-phase 5 μm C18 Luna column (Phenomenex), with a mobile phase consisting of 50% acetone (v/v) and 50% 20 mM aqueous formic acid (v/v), flowing at 0.6 mL/min. Absorbance was continuously monitored at 650 nm. Upper trace represents the product of an *Arabidopsis thaliana* HY2 assay (reaction incomplete due to leftover BV present in the chromatogram); Pre-1, reaction product of Pre-1 using BV as substrate. (C) Difference spectra of coupled phytochrome assembly assays employing apo-*Syn*Cph1 incubated with a PФB standard or the product of the Pre-2 reaction (apo-*Syn*Cph1 + Pre-2 prod.). Spectra were recorded after incubation for 3 min with red light (636 nm – Pfr spectrum) and following incubation for 3 min with far-red light (730 nm – Pr spectrum). Difference spectra were calculated by the subtraction of the Pfr from the Pr spectrum. The apo-*Syn*Cph1 + PФB difference spectrum (dashed line) was smoothed applying a 10 pt. Savitzy-Golay filter. The apo-*Syn*Cph1 + Pre-2 prod. difference spectrum (solid line) was smoothed applying a 20 pt. Savitzky-Golay filter and the signal was 3x enhanced.

**Figure 5.**
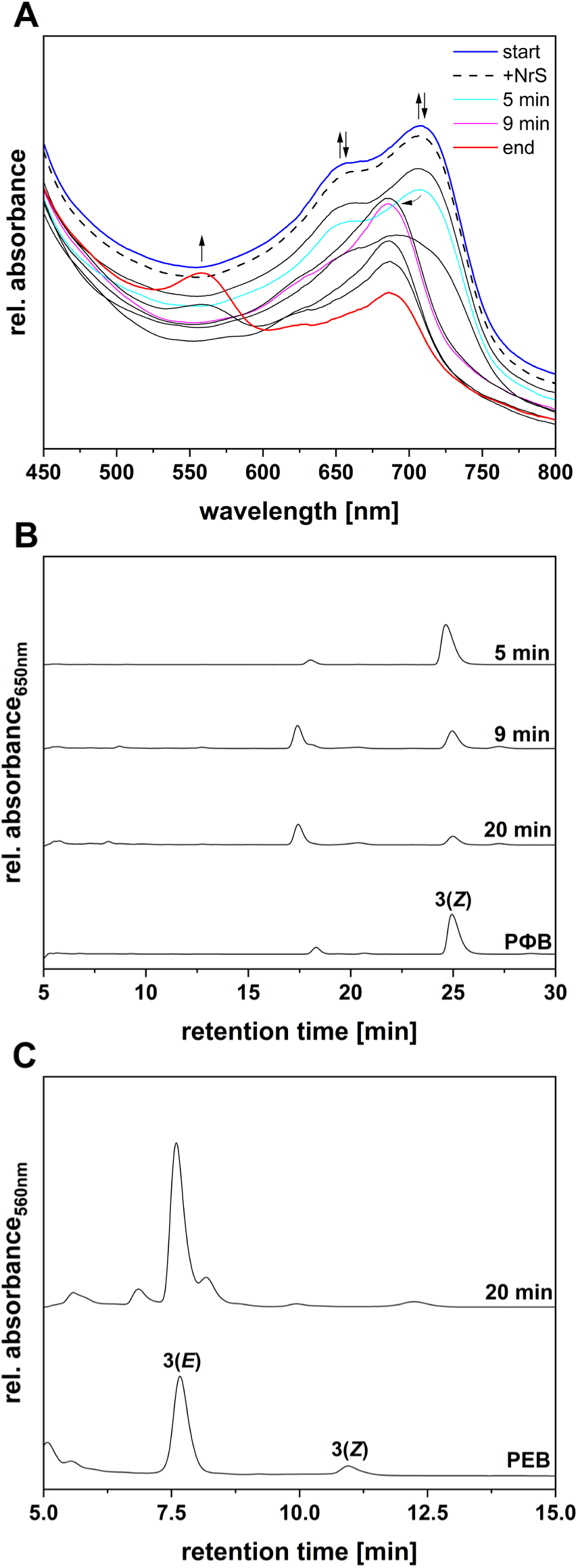
Investigation of the activity of recombinant Pre-3 and identification of reaction products. (A) Time-dependent spectra of an anaerobic bilin reductase activity assay using recombinant Pre-3 and BV as the substrate. The total reaction time was 20 minutes and spectra were recorded at 30 seconds intervals, although, for clarity, only informative spectra are shown. The arrows follow the absorbance changes during the reaction. The blue spectrum corresponds to the “binding spectrum”, recorded upon incubation of BV with Pre-3. The dashed spectrum represents the first recorded after initiating the reaction with the NrS. The black spectra indicate the most relevant ones recorded during the assay, while the final spectrum is red. An 40-points Savitzky-Golay filter was applied to smooth the curves. (B) HPLC analysis of the reaction products (5 min, 9 min, 20 min). The products were separated on a reversed-phase 5 μm C18 Luna column (Phenomenex), with a mobile phase consisting of 50% acetone (v/v) and 50% 20 mM aqueous formic acid (v/v), flowing at 0.6 mL/min. Absorbance was continuously monitored at 650 nm. PΦB, phytochromobilin standard (obtained from an *Arabidopsis thaliana* HY2 assay). (C) HPLC analysis of the 20 min reaction product. The products were separated on a reversed-phase 5 μm C18 Luna column (Phenomenex), with a mobile phase consisting of 50% acetone (v/v) and 50% 20 mM aqueous formic acid (v/v), flowing at 0.6 mL/min. Absorbance was continuously monitored at 560 nm. PEB, phycoerythrobilin standard.

### Diverse roles for catalytic residues in pre-PcyA active sites

To better understand the behavior of pre-PcyA proteins, we revisited potential catalytic residues corresponding to Glu76, His88, and Asp105 in PcyA (Figs. S1 and 2C). Glu76 is conserved in known PcyA proteins and is essential for reduction of the 18-vinyl moiety of BV by PcyA (Tu et al., 2007). This residue is conserved neither in PcyX members of the AX clade, which have an Asp at this position and synthesize PEB instead of PCB (Ledermann et al., 2016, 2018), nor in pre-PcyA proteins, which do not have a conserved residue at this position (Figs. S1 and 2C). His88 is conserved in PcyA and PcyX but not in other FDBRs; pre-1 enzymes have some variation at this position, whereas Pre-2 and Pre-3 have conserved Asn and Thr, respectively. Asp105 is conserved in PcyA, PcyX, Pre-2, and Pre-3 but typically corresponds to Glu in Pre-1. The D105N variant of PcyA results in a long-lived biliverdin radical, permitting detailed characterization of a normally unstable reaction intermediate (Tu et al., 2007; Stoll et al., 2009; Kohler et al., 2010). Hence, pre-PcyA proteins apparently have an equivalent acidic residue to Asp105 of PcyA, making this residue an interesting target for site-directed mutagenesis.

We extended these findings by comparing AlphaFold models of the three pre-PcyA proteins characterized in this study to the experimental crystal structure of *Syn*PcyA (PDB accession code: 2D1E) with bound BV substrate (Fig. 6). Results for CAP_1520 and MBL9008304 were similar to previous analysis using both AlphaFold predicted structures and conventional homology models (Rockwell et al., 2023), confirming that the conserved Asp of Pre-2 and Pre3 should be similarly positioned to Asp105 in PcyA. The predicted model for MCC5789364 (Pre-1) again placed the Pre-1 Glu residue in a similar position to Asp105, reinforcing the hypotheses drawn from the sequence alignment (Figs. 2C and 6A-C). Despite the ability of Pre-3 to reduce PΦB to PEB, its active site remains remarkably similar to that of *Syn*PcyA (Fig. 6C). We therefore considered alternative candidate catalytic residues. The only other FDBRs known to produce PCB from BV are HY2 proteins from streptophyte algae (Rockwell & Lagarias, 2017; Frascogna et al., 2023). Remarkably, an Asn-to-Asp substitution in one such protein at the position equivalent to His88 in PcyA resulted in synthesis of PEB rather than PCB (Frascogna et al., 2023), illustrating the extent to which subtle changes can produce major, surprising effects in FDBRs. Algal HY2 proteins also lack Glu76, but Asp residues equivalent to Asp105 of PcyA and Asp256 of Arabidopsis HY2 are critical for catalysis in *Kfla*HY2 (Frascogna et al., 2023). However, pre-3 proteins typically have basic residues at the latter position, such as Arg215 in MBL9008304 (Figs. S1 and 2C). However, the model structure for this protein placed a nearby residue, Glu217, close to the BV substrate and hence provided a candidate residue specific for reduction of the 15,16-double bond and PEB formation (Fig. 6C). We therefore examined four variant proteins: Pre-1_E91Q, Pre-2_ D91N, and Pre-3_D96N all replaced the conserved acidic residue with approximately isosteric neutral residues, whereas Pre-3_E217Q introduced a similar substitution at the proposed proton donor for 15,16 reduction. The activity of the recombinant enzymes was investigated *in vitro* via FDBR assays and subsequent HPLC analyses.

**Figure 6.**
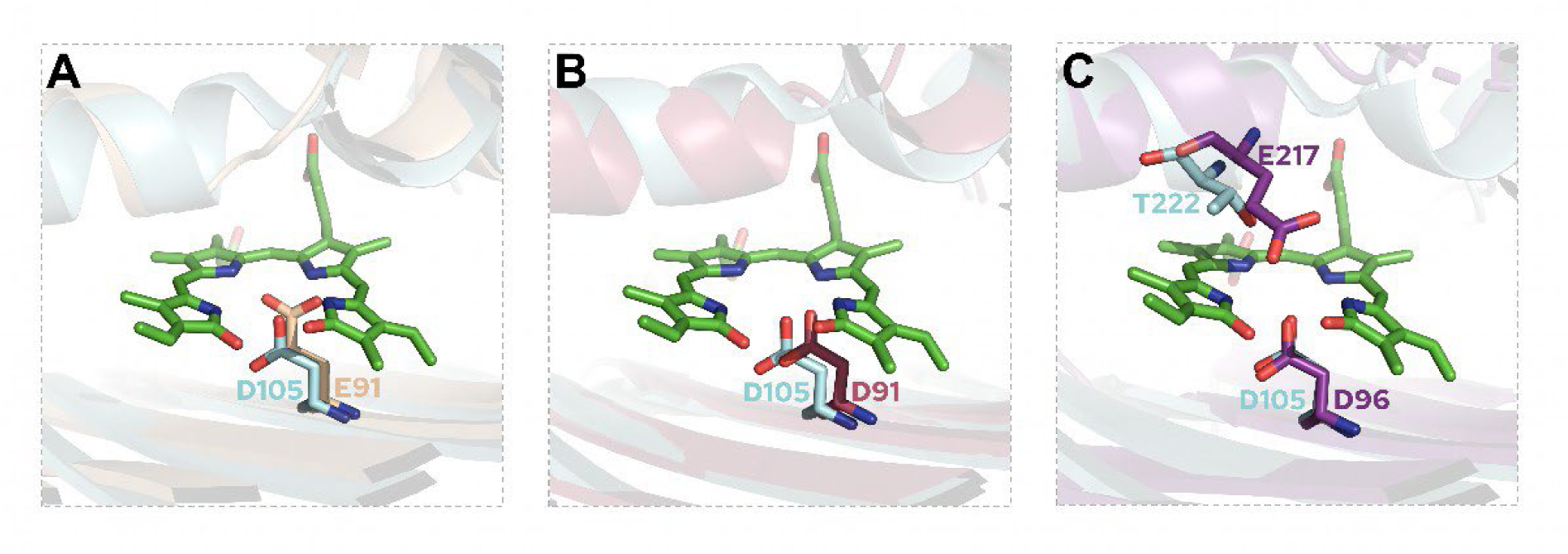
Structural modeling of pre-PcyA active sites and candidate catalytic residues. Structural alignments are shown for pre-PcyAs characterized in this study superimposed on the experimental structure of *Synechocystis* PcyA with bound BV (PDB accession 2D1E). Model structures of all pre-PcyAs were obtained using AlphaFold2 (Jumper et al., 2021; Mirdita et al., 2022), available at (https://colab.research.google.com/github/sokrypton/ColabFold/blob/main/AlphaFold2.ipynb). PyMOL (DeLano, 2020) was used to construct the superpositions. (A) Overlay of the active sites of MCC5789364 (Pre-1) and PcyA. The model structure of MCC5789364 is displayed as transparent wheat-colored cartoon. The crystal structure of PcyA is displayed as transparent aquamarine-colored cartoon. BV is represented as green sticks. Key amino acid residues are displayed as labeled sticks and colored according to the corresponding protein structure. (B) Overlay of the active sites of CAP_1520 (Pre-2) and PcyA. The model structure of CAP_1520 is displayed as transparent raspberry-colored cartoon. PcyA and BV are displayed as in panel A. Key amino acid residues are displayed as labeled sticks and colored by structure. (C) Overlay of the active sites of MBL9008304 (Pre-3) and PcyA. The model structure of MBL9008304 is displayed as transparent grape-colored cartoon. PcyA and BV are displayed as in panel A. Key amino acid residues are displayed as labeled sticks and colored by structure.

The Pre-1_E91Q variant was able to bind BV and use it as a substrate: initial formation of a peak at ∼ 725 nm was followed by appearance of a final product absorbing at ∼ 640 nm (Fig. 7A). HPLC analysis revealed an incomplete reaction (due to the presence of remaining BV) and the formation of mainly 3(Z)-PΦB, with traces of 3(*E*)-PΦB (Fig. 7B). This variant thus can still reduce BV, but the PΦB reaction product adopts the typical 3(*Z*) stereochemistry seen for other FDBRs. The equivalent D91N variant of Pre-2 was also able to bind BV and convert it to a final product absorbing at ∼ 645 nm (Fig. 7C). HPLC analysis revealed that this variant does not result in altered stereochemistry, with reduction of BV to 3(*E*)-PΦB (Fig. 7D) equivalent to that seen in the parent enzyme (Fig. 4B).

**Figure 7.**
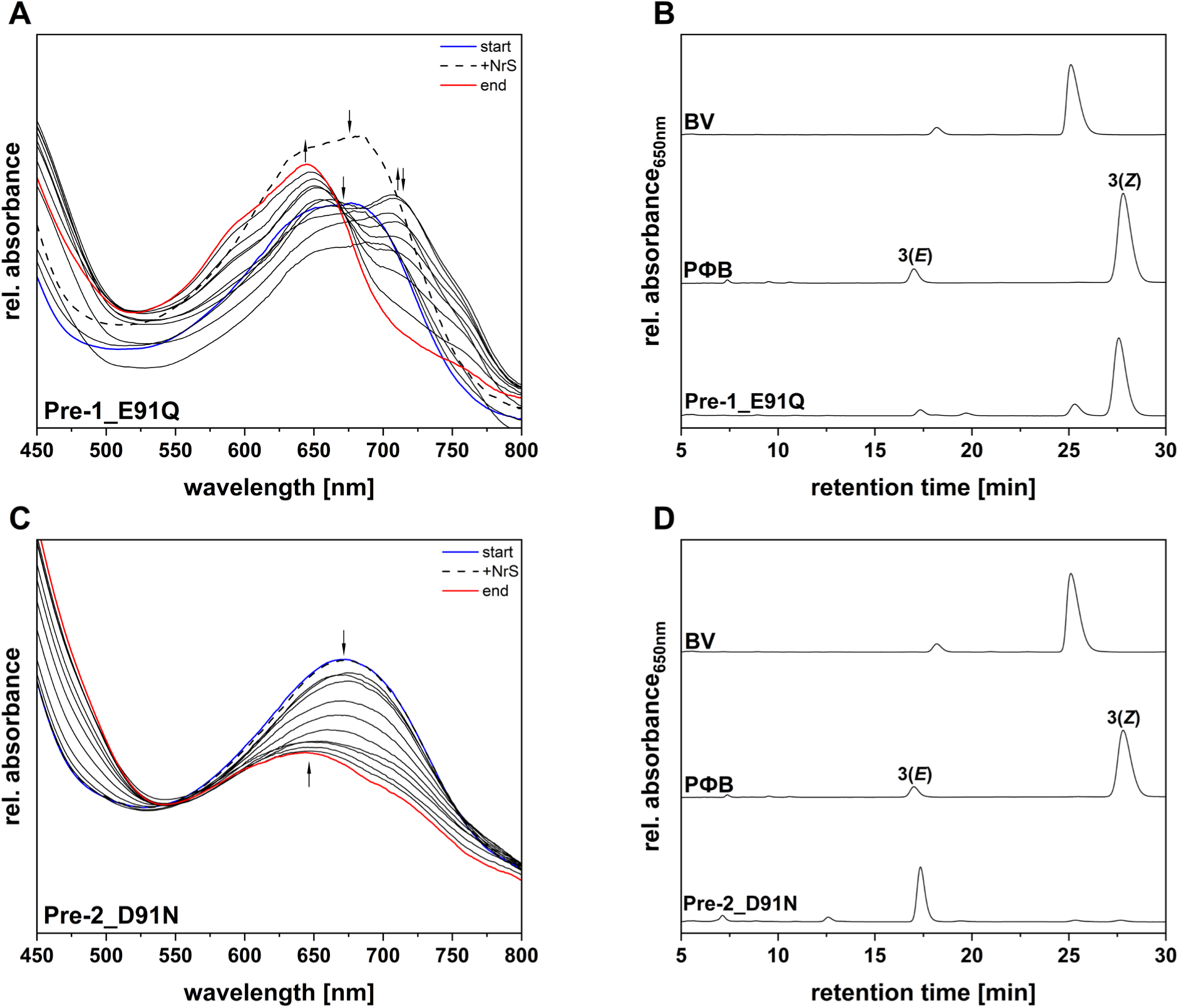
Investigation of the activity of recombinant Pre-1_and Pre-2 variants and identification of reaction products. (A) Time-dependent spectra of an anaerobic bilin reductase activity assay using recombinant Pre-1_E91Q and BV as the substrate. The total reaction time was 20 minutes and spectra were recorded at 30 seconds intervals, although, for clarity, only informative spectra are shown. The arrows follow the absorbance changes during the reaction. The blue spectrum corresponds to the “binding spectrum”, recorded upon incubation of BV with Pre-1_E91Q. The dashed spectrum represents the first recorded after initiating the reaction with the NrS. The black spectra indicate the most relevant ones recorded during the assay, while the final spectrum is red. A 50-points Savitzky-Golay filter was applied to smooth the curves. (B) HPLC analysis of the reaction products (Pre-1_E91Q). The products were separated on a reversed-phase 5 μm C18 Luna column (Phenomenex), with a mobile phase consisting of 50% acetone (v/v) and 50% 20 mM aqueous formic acid (v/v), flowing at 0.6 mL/min. Absorbance was continuously monitored at 650 nm. BV, biliverdin IXα standard; PΦB, phytochromobilin standard (obtained from an *Arabidopsis thaliana* HY2 assay). (C) Time-dependent spectra of an anaerobic bilin reductase activity assay using recombinant Pre-2_D91N and BV as the substrate. The total reaction time was 30 minutes and spectra were recorded at 30 seconds intervals, although, for clarity, only informative spectra are shown. The arrows follow the absorbance changes during the reaction. The blue spectrum corresponds to the “binding spectrum”, recorded upon incubation of BV with Pre-2_D91N. The dashed spectrum represents the first recorded after initiating the reaction with the NrS. The black spectra indicate the most relevant ones recorded during the assay, while the final spectrum is red. A 50-points Savitzky-Golay filter was applied to smooth the curves. (D) HPLC analysis of the reaction products (Pre-2_D91N). The products were separated on a reversed-phase 5 μm C18 Luna column (Phenomenex), with a mobile phase consisting of 50% acetone (v/v) and 50% 20 mM aqueous formic acid (v/v), flowing at 0.6 mL/min. Absorbance was continuously monitored at 650 nm. BV, biliverdin IXα standard; PΦB, phytochromobilin standard (obtained from an *Arabidopsis thaliana* HY2 assay).

Surprisingly, both Pre-3 variants exhibited a yellow-green coloration upon purification, and their absorbance and fluorescence spectra suggested the presence of potential porphyrin compounds. We assayed activity for these variants despite a high spectroscopic background. The activity assay performed for Pre-3_D96N revealed that it still potentially performed some kind of BV reduction, indicated by the presence of a double absorbance at ∼ 590 nm and ∼ 680 nm (Fig. 8A). HPLC analysis revealed production of small amounts of both C3 isomers of PΦB (Fig. 8B). We also noted the presence of a species eluting at ∼ 17 min which could not be assigned to any bilin. Pre-3_E217Q exhibited clearer activity in the spectroscopic assay (Fig. 8C). In this case, the reduction of BV to PΦB was almost complete (Fig. 8D). Moreover, the unknown species recorded in the previous case was also found.

**Figure 8.**
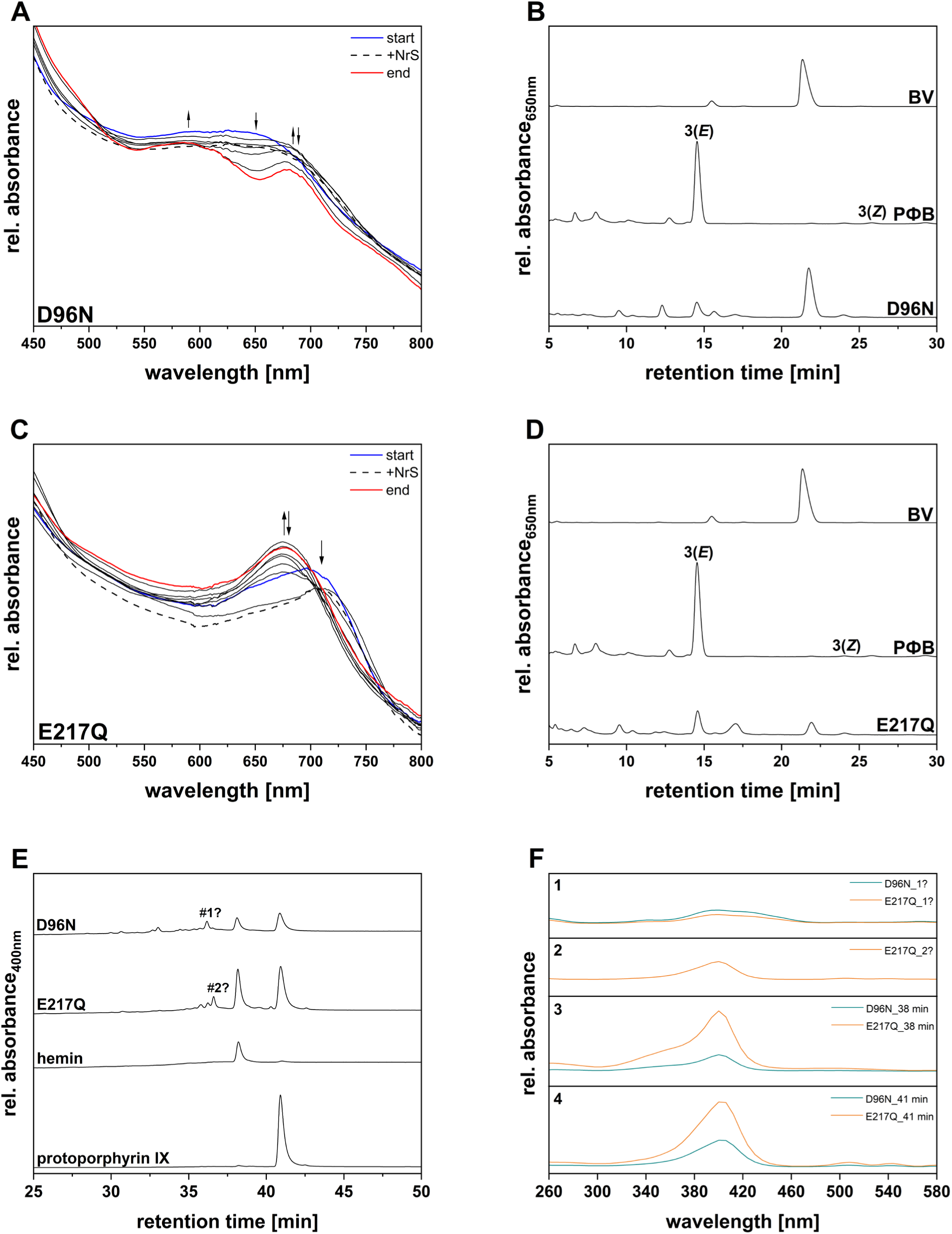
Investigation of the activity of recombinant Pre-3 variants, identification of reaction products and extracted tetrapyrroles. (A) Time-dependent spectra of an anaerobic bilin reductase activity assay using recombinant Pre-3_D96N and BV as the substrate. The total reaction time was 30 minutes and spectra were recorded at 30 seconds intervals, although, for clarity, only informative spectra are shown. The arrows follow the absorbance changes during the reaction. The green spectrum corresponds to the “binding spectrum”, recorded upon incubation of BV with Pre-3_D96N. The dashed spectrum represents the first recorded after initiating the reaction with the NrS. The black spectra indicate the most relevant ones recorded during the assay, while the final spectrum is red. A 70-points Savitzky-Golay filter was applied to smooth the curves. (B) HPLC analysis of the reaction products (Pre-3_D96N). The products were separated on a reversed-phase 5 μm C18 Luna column (Phenomenex), with a mobile phase consisting of 50% acetone (v/v) and 50% 20 mM aqueous formic acid (v/v), flowing at 0.6 mL/min. Absorbance was continuously monitored at 650 nm. BV, biliverdin IXα standard; PΦB, phytochromobilin standard from Pre-1 assay (20 min) using BV as substrate. (C) Time-dependent spectra of an anaerobic bilin reductase activity assay using recombinant Pre-3_E217Q and BV as the substrate. The total reaction time was 20 minutes and spectra were recorded at 30 seconds intervals, although, for clarity, only informative spectra are shown. The arrows follow the absorbance changes during the reaction. The green spectrum corresponds to the “binding spectrum”, recorded upon incubation of BV with Pre-3_E217Q. The dashed spectrum represents the first recorded after initiating the reaction with the NrS. The black spectra indicate the most relevant ones recorded during the assay, while the final spectrum is red. A 100-points Savitzky-Golay filter was applied to smooth the curves. (D) HPLC analysis of the reaction products (Pre-3_E217Q). The products were separated on a reversed-phase 5 μm C18 Luna column (Phenomenex), with a mobile phase consisting of 50% acetone (v/v) and 50% 20 mM aqueous formic acid (v/v), flowing at 0.6 mL/min. Absorbance was continuously monitored at 650 nm. BV, biliverdin IXα standard; PΦB, phytochromobilin standard from pre-PcyA assay using BV as substrate. (E) HPLC analysis of the tetrapyrroles extracted from Pre-3_D96N and Pre-3_E217Q. The products were separated on a reversed-phase 5 μm Equisil BDS C18 column (Dr. Maisch HPLC GmbH), using a gradient of 9% acetonitrile in 1 M ammonium acetate pH 5.2 and 9% acetonitrile in methanol, flowing at 0.5 mL/min (Kühner et al., 2014). Absorbance was continuously monitored at 400 nm. Hemin and Protoporphyrin IX were used as standards. (F) Absorbance spectra of HPLC-detected compounds in Figure 8A. Panel 1 shows the absorbance of the compound denoted as “#1?”. Panel 2 shows the absorbance of the compound denoted as “#2?”. Panel 3 shows the absorbance of the compound eluting at ∼ 38 min. Panel 4 shows the absorbance of the compound eluting at ∼ 41 min. Spectra referred to the products isolated from Pre-3_D96N are depicted in dark cyan, whereas the ones isolated from Pre-3_E217Q are depicted in orange.

In order to examine potential porphyrins co-purifying with the Pre-3 variants, we extracted pigments using ethyl acetate (Kühner et al., 2014). Based on subsequent HPLC analysis (Fig. 8E), Pre-3_D96N had heme and protoporphyrin IX bound, in addition to a third unknown pigment (Fig. 8E). The same species were identified in the Pre-3_E217Q variant, along with an additional peak (Fig. 8E-F). These data suggest that the introduced active site variations generated Pre-3 active sites with higher affinity for cyclic tetrapyrroles along with prevention of PEB formation.

## Discussion

The recent discovery of FDBR-like sequences in heterotrophic, non-photosynthetic bacteria offers a valuable opportunity to investigate the evolutionary development of this enzyme family and its functional diversification. These sequences cluster in three lineages: pre-1, pre-2 and pre-3 (Fig. S1; Rockwell et al., 2023; Frascogna, Rockwell et al., 2026). Previous work (Rockwell et al., 2023) demonstrated FDBR activity for representative Pre-2 and Pre-3 proteins using an indirect co-expression assay, with little activity seen for Pre-1. We here show that the PEB isomer produced by Pre-3 in that assay is distinct from that produced by the distantly related FDBR PebS (Fig. 1 and Table 1), highlighting the need to characterize pre-PcyA proteins *in vitro* for detailed understanding.

One representative sequence from each was selected for biochemical analysis. Due to the poor activity observed with the previously selected Pre-1 (POZ53557), we identified an alternative Pre-1 candidate (MCC5789364; Fig. 2), to more clearly define Pre-1 activity. These three FDBRs were then characterized after recombinant expression and purification. The results unveiled two previously unreported scenarios for FDBRs. First, pre-PcyA proteins carry out the 2-electron reduction of BV to 3(*E*)-PΦB (Figs. 3–5). Previously characterized FDBRs instead favor the production of 3(*Z*) phytobilin isomers (Frankenberg & Lagarias, 2003; Tu et al., 2008; Ledermann et al., 2016). Interestingly, the latter was also identified as a potential precursor to 3(*E*)-PCB and PΦB in older *in vivo* studies (Beale & Cornejo, 1984; Terry et al., 1995; J. L. Weller et al., 1996). The *E* isomer is thermodynamically more stable than the *Z* (Weller & Gossauer, 1980); indeed, both ethylidene isomers of PCB can be isolated from phycocyanin by chemical cleavage, but the 3(*Z*) isomer converts spontaneously into 3(*E*) under high-temperature methanolysis or ethanolysis (Fu et al., 1979; Cornejo, 1992; Terry et al., 1993; Roda-Serrat et al., 2018; Kamo et al., 2021; Wilkinson et al., 2023). This heat-induced reaction was also suspected to cause partial isomerization from 3(*Z*) to 3(*E*) in early FDBR research, likely during SpeedVac-mediated concentration of bilins (Frankenberg et al., 2001). In this study, however, this step was replaced with sublimation using a lyophilizer, thereby preventing potential heat-induced isomerization. This indicates that 3(*E*)-PΦB here cannot be attributed to thermal artifacts, making pre-PcyA proteins the first known FDBRs to preferentially synthesize 3(*E*) phytobilins (Fig. 9).

**Figure 9.**
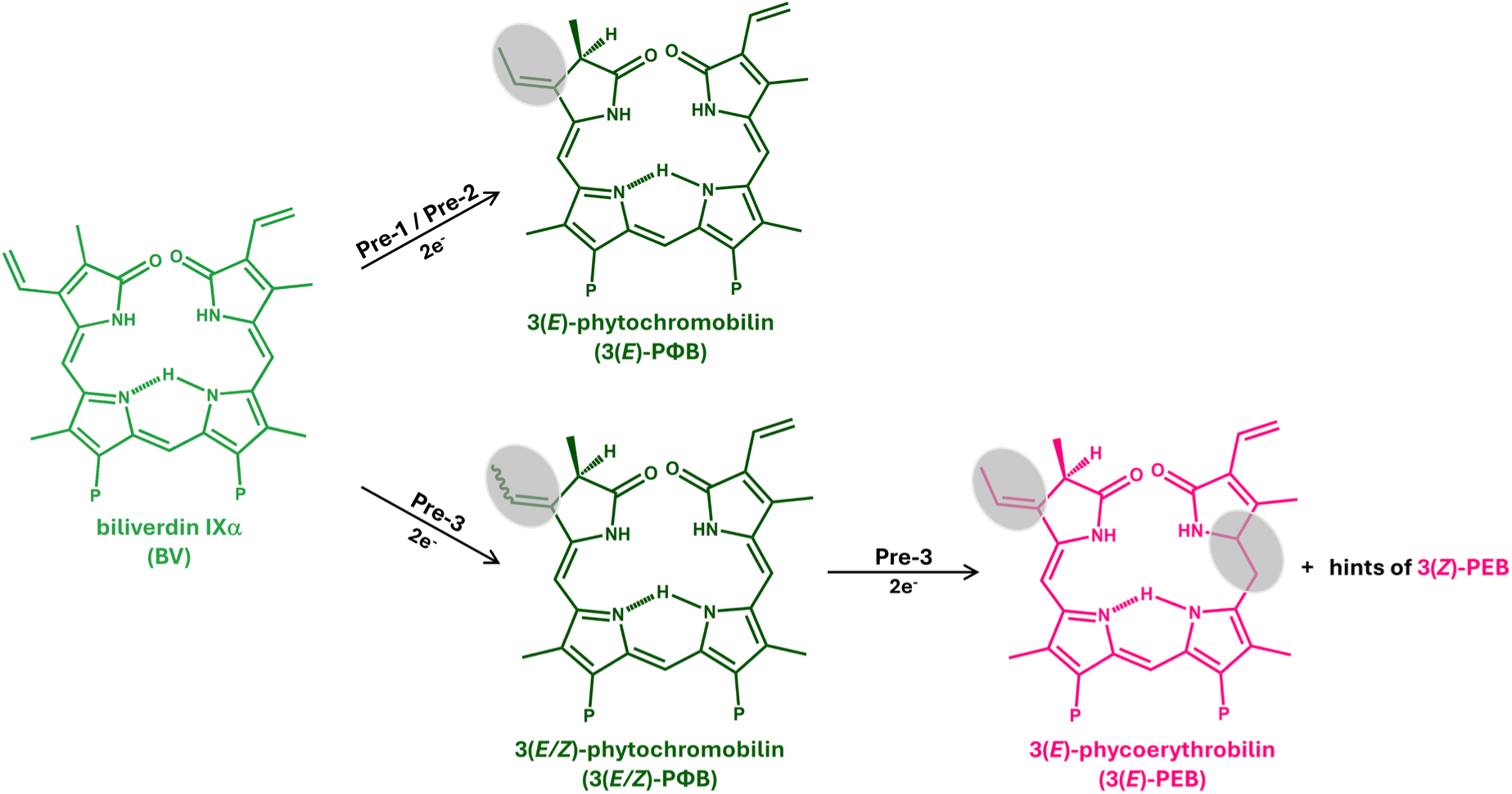
Reactions catalyzed by the pre-PcyAs. Pre-1 and Pre-2 catalyze the two-electron reduction of BV to 3(*E*)-PΦB. Pre-3 catalyzes the four-electrons reduction of BV to PEB via the intermediate PΦB. BV reduction sites are highlighted in gray. “P” in the bilin structures indicates the propionate side chains.

The second finding from *in vitro* characterization of wild-type enzymes is the confirmation of PEB formation by Pre-3. After converting the A-ring diene system of BV to PΦB, Pre-3 further reduces the C15=C16 double bond, yielding 3(*E*)-PEB. Reduction of PΦB to PEB has previously only been observed for PebA, which can reduce mixture of 3(Z)/3(*E*)-PΦB to *3*(*Z*)*/3*(*E*)-PEB (Dammeyer & Frankenberg-Dinkel, 2006). However, BV is the physiological substrate for PebA, so this observation is an artifact arising from an intrinsic lack of PebA specificity. In contrast, the activity of Pre-3 was recorded with the *bona fide* substrate, BV: PEB was also formed from PΦB in the *in vivo* co-expression assay incorporating HO to synthesize BV, and the open reading frame for the pre-3 MBL9008304 is located within a cluster that also includes both a bilin-binding acceptor protein (the BBAG MBL9008307) and HO (MBL9008303; Rockwell et al., 2023). However, the formation of 3(*E*)-PEB by Pre-3 explains the distinct populations of PEB apparently produced by this enzyme and by PebS (Fig. 1): the discrepancy arises because the BBAG MBL9008307 can incorporate either 3(*E*) or 3(*Z*) phytobilins, but the two isomers lead to populations with slightly different spectral properties.

We next compared predicted AlphaFold structures for all three pre-PcyA proteins to the experimental structure for *Synechocystis* PcyA with bound BV (Fig. 6) as a means of identifying active-site residues implicated in pre-PcyA catalysis and regiospecificity. We first targeted a conserved acidic residue for site-directed mutagenesis. This residue corresponds to the critical Asp105 of PcyA (Table 2), with Asp-to-Asn substitutions in that enzyme resulting in formation of long-lived radical intermediates and poor overall catalysis (Tu et al., 2007). Similar results were not obtained with pre-PcyA enzymes. In Pre-2, the D91N variant retained the activity of the wild-type protein. The equivalent E91Q variant in Pre-1 resulted in inversion of stereochemistry in the reaction product, yielding 3(*Z*)-PΦB. In Pre-3, the equivalent D96N did exhibit greatly reduced activity; however, far-red-absorbing radical species were not accumulated during the reaction, and trace amounts of product were formed. We also examined the behavior of the E217Q variant of Pre-3; based on the predicted position of this residue in the AlphaFold structure, it seemed a possible proton donor for reduction of the 15,16-double bond. Indeed, PEB formation was not observed in this variant. Instead, it exhibited inefficient synthesis of 3(*E*)-PΦB. Hence, this variant was still able to carry out a complete 2-electron reduction of BV, in contrast to the failure of D105N PcyA to carry out reduction of BV to 18^1^,18^2^-DHBV. These substitutions in Pre-3 thus provide further support for the formation of PEB exclusively via PΦB: the substitution of Glu217 still leads to the production of PΦB intermediate, meaning that Glu217 is important for D-ring reduction but not for overall activity, whereas substitution of Asp96 results in greatly reduced activity.

Both Pre-3 variants also co-purified with endogenous pigments after recombinant expression in *E. coli*, resulting in visibly yellow-green protein preparations. Extracting these preparations with ethyl acetate allowed identification of protoporphyrin IX and heme, along with two unknown tetrapyrroles. This result complicates analysis of the effects of the substitutions, because it implies a shift in ligand specificity from linear to cyclic tetrapyrroles. The presence of other tetrapyrroles in the protein sample also could interfere with the activity assay, leading to high spectral background and potentially also hindering the overall activity of the enzyme by competing with the binding of BV substrate.

The ability of these variants to bind cyclic tetrapyrroles is noteworthy, especially in light of observations made at the time of the crystallization of the first FDBR, PcyA from *Synechocystis* sp. PCC 6803 (Hagiwara et al, 2006). At that time, an unexpected structural similarity to oxygen-dependent coproporphyrinogen III oxidases (CPOs) from *Leishmania* and *Saccharomyces cerevisiae* was found despite a lack of statistically significant sequence similarity. Indeed, the overall fold is highly conserved between these otherwise functionally unrelated enzymes (Fig. 10A). There is thus a precedent for binding of cyclic tetrapyrroles to this overall fold, as seen in the Pre-3 variants. However, the FDBR and CPO active sites are on opposite sides of the central β-sheet in this fold (Figure 11), and the CPO reaction is an oxidative decarboxylation rather than a reduction (Fig. 10B). Thus, the observed shift in substrate specificity likely arises from a novel structural adaptation rather than a convergent reuse of the CPO catalytic architecture. Furthermore, oxygen-dependent CPOs function as dimers, whereas FDBRs are monomeric enzymes (Phillips et al., 2004; Lee et al., 2005; Overkamp & Frankenberg-Dinkel, 2013).

**Figure 10.**
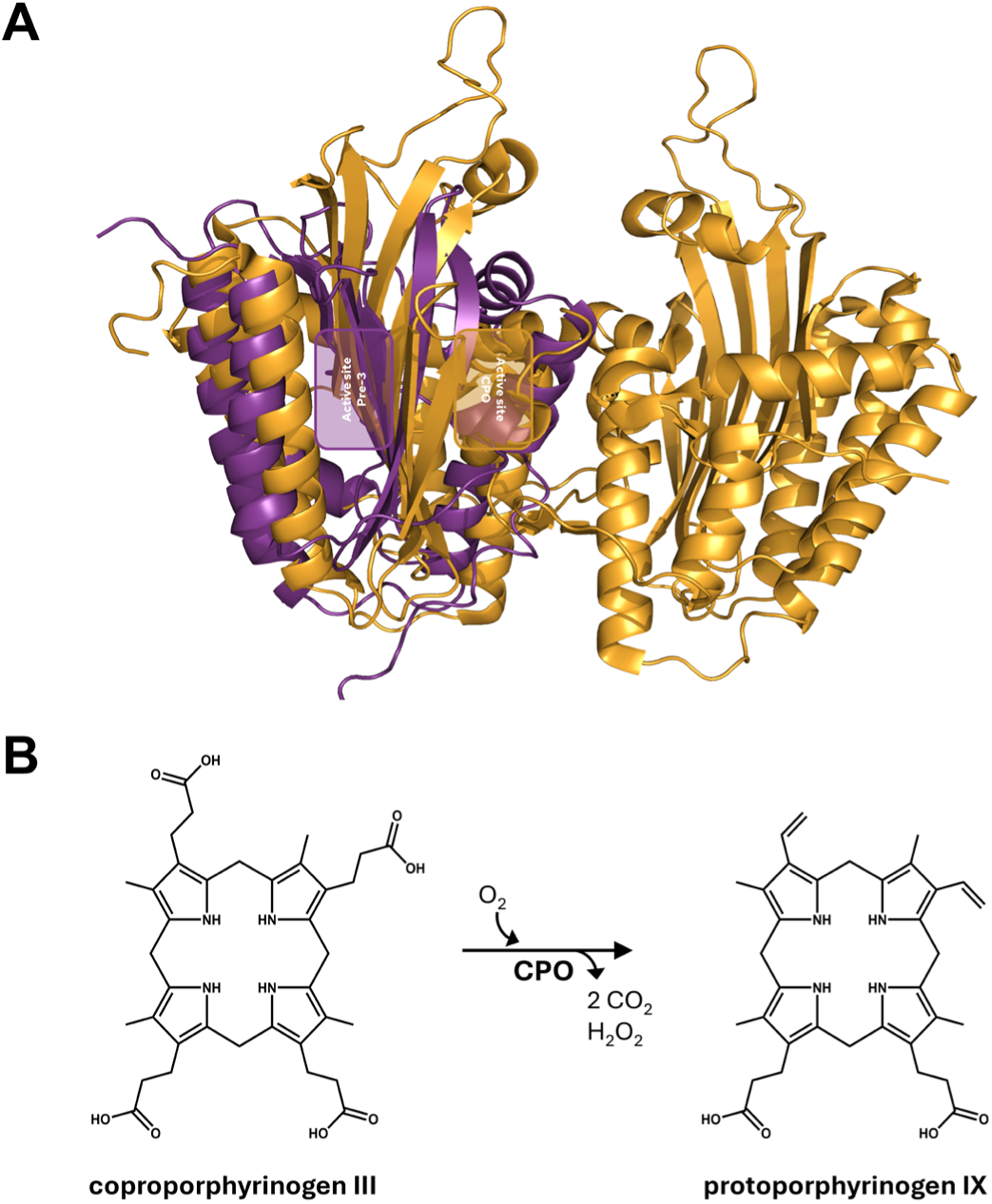
Pre-3 and yeast coproporphyrinogen III oxidase. (A) Overlay of Pre-3 with oxygen-dependent coproporphyrinogen III oxidase (CPO) from Saccharomyces cerevisiae (PDB accession code: 1TKL). PyMOL (DeLano, 2020) was used to construct the topology-based alignment using the Combinatorial Extension algorithm (CEalign function). The model structure of Pre-3 was obtained using AlphaFold2 (https://colab.research.google.com/github/sokrypton/ColabFold/blob/main/AlphaFold2.ipynb) (Jumper et al., 2021; Mirdita et al., 2022). The model structure of Pre-3 is displayed as violet purple-colored cartoon. The crystal structure of the CPO is displayed as bright orange-colored cartoon. Active sites are highlighted. (B) The reaction catalyzed by CPO is shown.

Overall, our work reinforces the structural plasticity of the FDBR scaffold and provides new insight into the difficulty in rationally controlling regiospecificity by site-directed mutagenesis (Dammeyer et al., 2008; Ledermann et al., 2018; Frascogna et al., 2023; Miyake et al., 2025). Equivalent substitutions in Pre-2 and Pre-3 have radically different effects: in Pre-2, D91N has no effect, but the D96N variant of Pre-3 has higher affinity for cyclic tetrapyrroles and reduced FDBR activity. E91Q Pre-1 retains its normal 2-electron reduction but results in inverted stereochemistry for the reaction product. We propose that these unpredictable and varied effects arise at least in part via effects on substrate positioning relative to other potential catalytic residues. Thus, for example, E91Q would remove a proton donor from the Pre-1 active site by analogy to the effects of D105N in PcyA. However, it might also shift the position of the BV substrate and make an alternate proton donor more able to facilitate catalysis. The location of this alternate donor would then determine the final stereochemistry of the product. Conversion between 2-electron and 4-electron reactions in FDBRs might also be complicated by similar factors; without proton donation, radical intermediates do not proceed to yield stable products. Within the overall evolution of FDBRs (Fig. S1), conversion between 2-electron and 4-electron reduction has occurred repeatedly. The three known FDBR clades are PcyA/PcyX, PebA/PebS/PUBS, and PebB/HY2. All three are now known to carry out either 2-electron or 4-electron reductions: pre-PcyA proteins belong to the PcyA/PcyX clade and themselves can carry out either reaction, whereas PebA is a 2-electron reductase and PebS is a 4-electron reductase. HY2 enzymes from different organisms vary, with algal enzymes carrying out 4-electron reductions and those from land plants instead carrying out 2-electron reductions. PebS is an early branch within the PebA/PebS/PUBS clade, which is the first FDBR clade to branch relative to the CPO outgroup (Fig. S1). This indicates that 4-electron reductions appeared fairly early in FDBR evolution. To unravel the early steps in the evolution of modern bilin biosynthesis, it will thus be interesting to see whether earlier “pre-PebA” or “archae-FDBR” enzymes can be identified.

## Materials and Methods

### Phylogenetic analysis

Our recent phylogeny of FDBRs (Frascogna, Rockwell et al., 2026) was updated by inclusion of additional sequences from kelps (brown algae, Phaeophyceae). These sequences were chosen because they were only distantly related to pre-PcyA sequences, so changes in the resulting tree would be expected to arise from intrinsic instability to facilitate identification of robustly assigned pre-1 sequences. We then constructed a multiple sequence alignment in MAFFT v7.450 (Katoh & Standley, 2014) using the E-INS-i algorithm (equivalent to the command-line settings --genafpair --maxiterate 16 --clustalout --reorder). Alignment positions having gaps at ≥5% of the total sequences were removed using an in-house script to yield a final alignment having 350 sequences and 166 characters. The final alignment was then used to infer a maximum likelihood phylogeny in IQ-Tree 3.0.1 (May 5 2025 build; https://iqtree.github.io/) using automatic model selection with ModelFinder (Kalyaanamoorthy et al., 2017) (command-line settings: -msub nuclear -alrt 1000 -B 1000 -T AUTO), with supports evaluated using UFBoot and SH-aLRT (Guindon et al., 2010; Hoang et al., 2018). Trees were visualized and processed in TreeViewer (Bianchini & Sánchez-Baracaldo, 2024). Active-site residues were assessed using the original sequence alignment prior to gap removal, using the numbering of PcyA from *Synechocystis* sp. PCC 6803 (Hagiwara et al., 2006) and HY2 from *Arabidopsis thaliana* (Kohchi et al., 2001).

### Chemicals

All chemicals used in this study were ACS grade or better. Biliverdin IXα was purchased from Frontier Specialty Chemicals (Newark, Delaware, USA). Ferredoxin (PetF) and Ferredoxin-NADP^+^ reductase (FNR or PetH) were recombinantly produced and purified as described elsewhere (Busch et al., 2011; Dammeyer et al., 2008). The remaining components for the activity assays were purchased from Sigma-Aldrich (Taufkirchen, Germany). HPLC-grade acetone, formic acid and acetonitrile were purchased from VWR Chemicals (Darmstadt, Germany).

### Plasmids and primers

Plasmids for co-expression of bilin binding proteins in the pET28(a) backbone with the pre-PcyA proteins CAP_1520 and MBL9008304 have been previously described (Rockwell et al., 2023). For co-expression with PebS, the same strategy was used by combining amino acids 2-233 of PSSM2_058 (Rockwell et al., 2023) with the published modifications (modified N-terminal sequence for better expression and introduction of an N-terminal FLAG tag). The resulting synthetic gene was obtained from Genscript (Piscataway, New Jersey, USA) after cloning into the BamHI and SpeI sites of plasmid Spam-545/alt (Rockwell et al., 2023). Other synthetic gene fragments (accession codes in Table 3) were purchased from Twist Bioscience (South San Francisco, California, USA). The plasmids in Table 3 were generated via Gibson assembly (Gibson et al., 2009), employing appropriate oligonucleotide primers. Protein variants were generated via site-directed mutagenesis.

**Table 3:** Plasmids used for expression and purification of pre-PcyA proteins.

| Plasmid | Features | Accession code |
| --- | --- | --- |
| pET28a_MCC5789364 | pET-28a(+) derivative carrying the reverse-translated gene sequence of putative FDBR MCC5789364 from *Opitutales* bacterium metagenome; N-terminal His<sub>6</sub>-tag | MCC5789364.1 (GenBank) |
| pET28a_CAP_1520 | pET-28a(+) derivative carrying the reverse-translated gene sequence of putative FDBR CAP_1520 from *Chondromyces apiculatus* DSM 436; N-terminal His<sub>6</sub>-tag | EYF06823 (GenBank) |
| pET28a_MBL9008304 | pET-28a(+) derivative carrying the reverse-translated gene sequence of putative FDBR MBL9008304 from *Myxococcales* bacterium metagenome; N-terminal His<sub>6</sub>-tag | MBL9008304 (GenBank) |
| pET28a_MCC5789364_E91Q | pET28a_MCC5789364 derivative obtained by site-directed mutagenesis |  |
| pET28a_CAP_1520_D91N | pET28a_CAP_1520 derivative obtained by site-directed mutagenesis |  |
| pET28a_MBL9008304_D96N | pET28a_MBL9008304 derivative obtained by site-directed mutagenesis |  |
| pET28a_MBL9008304_E217Q | pET28a_MBL9008304 derivative obtained by site-directed mutagenesis |  |

### Production and purification of recombinant proteins

For the production of recombinant His-tagged FDBRs, 2 L of LB medium supplemented with 50 µg/mL kanamycin was inoculated 1:100 with an overnight culture of *E. coli* BL21(DE3) carrying the pET28a derivatives. The cultures were grown at 37°C and 100 rpm (New Brunswick™ INNOVA^®^ 44) to an OD_600_ of 0.4 – 0.6. The temperature was decreased to 17°C and gene expression was induced by addition of 1 mM isopropyl-ß-thiogalactoside (IPTG). The cultures were incubated under shaking for 19 additional hours and harvested by centrifugation for 10 min at 17000 × g and 4°C (Sorvall LYNX 6000, Rotor F9). The pellet was resuspended in “His-Binding buffer” (20 mM sodium phosphate pH 7.4; 500 mM NaCl) in a ratio of 3 mL buffer/g wet cell weight. After the addition of DNaseI (AppliChem GmbH, Darmstadt, Germany) and lysozyme (Sigma-Aldrich), the suspension was kept on ice for 30 min. The cells were disrupted using a microfluidizer (LM 10 Microfluidizer, Microfluidics™; 3 cycles; 15000 psi) and centrifuged for 45 min at 50000 × g and 4°C (Sorvall LYNX 6000, Rotor T29). The crude extract was loaded onto a gravity flow column containing 2 mL of TALON^®^ Superflow™ resin (Cytiva, Freiburg im Breisgau, Germany). Following a washing step with 10 column volumes (CV) of “His-Binding buffer”, the elution was performed using 4 CV of “His-Elution buffer” (20 mM sodium phosphate pH 7.4; 500 mM NaCl; 500 mM imidazole). Elution fractions containing the target protein were pooled and dialyzed overnight against TES-KCl buffer (25 mM TES/KOH pH 7.5; 100 mM KCl; 10% glycerol [v/v]).

Co-expression and purification of BBAG MBL9008307 used a modification of the previously published procedure (Rockwell et al., 2023). 200 mL of LB pre-warmed to 37°C was supplemented with 1 mL of a freshly prepared, sterile filtered 0.1 M solution of δ-aminolevulinic acid hydrochloride (Frontier Specialty Chemicals), 80 µL kanamycin (50 mg/mL stock solution in water), and 134 µL chloramphenicol (30 mg/ml stock solution in 95% ethanol) and was then inoculated with 1 mL of an overnight culture of *E. coli* strain C41 carrying the desired plasmids. After growth to OD 0.8-1.0 at 37°C and 200 rpm, 3 mL of a freshly prepared, sterile filtered 0.5 M solution of IPTG (Teknova) was added. Culture temperature was reduced to 20°C, and the culture was shaken at 120 rpm for approximately 18 hours. Cells were then collected by centrifugation (10 min, 5000 × *g*, 20°C, Beckman JLA-16.250 rotor), and supernatant was discarded. The cell pellets were weighed and stored at -70°C to await purification. For purification, cell pellets were thawed in the dark. 6 mL NEBexpress lysis reagent (New England Biolabs), 60 µL 0.1 M CaCl_2_, 6 µL T4 lysozyme (New England Biolabs), and 6 µL micrococcal nuclease (New England Biolabs) were added per 1 g cell wet weight. Cells were then lysed by incubation in the dark for 20 minutes at room temperature with gentle agitation (100 rpm on an orbital shaker). Unlysed cells and cellular debris were removed by centrifugation (10 min, 20000 × *g*), and the supernatant was then loaded by gravity flow onto a Ni-NTA affinity column (Millipore His-Bind resin) pre-equilibrated in Buffer A/S (20 mM HEPES/NaOH, pH 7.8; 0.1 M NaCl; 10% glycerol [v/v]) at room temperature. The column was washed with ≥5 column volumes of Buffer A/S supplemented with 50 mM imidazole at 0.9 ml/min, followed by elution using Buffer A/S supplemented with 250 mM imidazole. Peak fractions were pooled, concentrated to ≤0.3 mL using microconcentrators (Amicon Ultra 0.5 mL, 10 kD cutoff), and applied to a Sephacryl S200HR gel filtration column (1.5x20 cm Kontes flex-column; approximate bed volume 30 ml) at room temperature pre-equilbrated with TKKG buffer (25 mM TES-KOH pH 7.8, 100 mM KCl, 10% (v/v) glycerol). Protein was then eluted in TKKG at 0.9 mL/min. Peak fractions were then pooled and characterized using absorption and fluorescence spectroscopy. Absorption spectra were acquired using a Cary60 spectrophotometer at 25°C (250-900 nm, sampled every 2 nm for 0.125 s). For acid denaturation, 100 µL protein samples were diluted in 1 mL of 7 M guanidinium chloride/1% HCl (v/v). Absorption spectra were then recorded as for native samples. Fluorescence spectra were acquired on a QM-6/2005SE fluorimeter equipped with red-enhanced photomultiplier tubes (Photon Technology International 814 Series). Some fluorescence spectra were acquired using a Schott glass 455 nm long-pass filter to suppress harmonics. MBL9008307 purified after co-expression with MBL9008304 using this procedure was spectroscopically indistinguishable from previously described material (Rockwell et al., 2023).

### Activity assay

The anaerobic bilin reductase activity assays were performed following the method outlined by Ledermann and colleagues with slight modifications (Ledermann et al., 2016). Recombinant ferredoxin PetF from the cyanophage P-SSM2 served as the electron donor at a final concentration of 1 µM. PetF was reduced using recombinant PetH from *Synechococcus* sp. PCC 7002 at a final concentration of 0.01 µM (Busch et al., 2011). The reaction was initiated by adding an NADPH regenerating system (NRS) consisting of 65 mM glucose-6-phosphate, 8.2 µM NADP+ and 11 U/ml glucose-6-phosphate dehydrogenase. Absorbance measurements were carried out using an Agilent 8453 spectrophotometer. The reaction was stopped by diluting the reaction mix 1:10 in ice-cold 0.1%_v/v_ TFA. Bilin products were isolated via solid phase extraction using Sep-Pak^®^ C18 Plus Light cartridges (Waters, Milford, Massachusetts, USA) and freeze-dried using an Alpha 2-4 LSC plus lyophilizer (Martin Christ GmbH, Osterode, Germany).

### Bilin identification

The isolated assay products were analyzed using an Agilent 1100 series chromatograph equipped with a Luna^®^ 5µm reversed phase C18-column (Phenomenex, Torrance, California, USA) and a diode-array detector. The mobile phase consisted of 50%_v/v_ acetone and 50%_v/v_ 20 mM formic acid, flowing at 0.6 mL/min. Reaction products were identified by comparing their retention times with known standards as well as full-spectrum analysis of the elution peaks.

*In vitro* chromophore assembly of the apo-phytochrome *Syn*Cph1 was carried out using the lysate of a pET_*cph1* overexpression after centrifugation and filtration. The assembly was tested employing 300 μL lysate incubated either with 40 μM standard bilin or with 4 μL of bilin solutions of unknown concentration for 30 min at room temperature, in the dark. Afterwards, the volume was adjusted to 500 μL with PBS. Absorbance spectra were recorded after incubation for 3 min with red light (636 nm – Pfr spectrum) and after incubation for 3 min with far-red light (730 nm for *Syn*Cph1 – Pr spectrum) using an Agilent 8453 spectrophotometer. Difference spectra were calculated by the subtraction of the Pfr from the Pr spectrum.

### Extraction of tetrapyrroles and HPLC analysis

For the extraction of protein-bound tetrapyrroles, 3 x 0.5 mL of protein solution were acidified each with 10 µL HCl (37%) followed by the addition of 0.7 mL of ethyl acetate. The solutions were vigorously mixed for ten minutes using a Disruptor Genie (VWR) and then centrifuged for 10 min at 4°C and 15.000 rpm in an Eppendorf 5427R centrifuge. 600 µL of the organic phase were transferred into a new 1.5 mL vial and concentrated to complete dryness. The dried pellets were stored at -20°C until use.

For HPLC analysis, the dried pellets were dissolved in 10 µL acetone/HCl (97.5/2.5). Three dissolved samples were combined and diluted with 100 µL of the HPLC start condition. The sample was acidified with 2.5 µL HCl (37%). After mixing and a 10 min centrifugation, 30 µL of the solution were applied to an Equisil BDS C18 column (250 x 4.6 mm, 5 µm) (Dr. Maisch GmbH, Ammerbuch-Entringen, Germany) attached to a JASCO HPLC 2000 series system (JASCO Deutschland GmbH, Pfungstadt, Germany). Separation of tetrapyrroles was performed as described previously (Kühner et al., 2014).

## Author contributions

FF, NCR, and NFD designed research; FF, NCR, and GL performed research; FF, NCR, GL and NFD wrote the paper. All authors read and approved the final version of the manuscript.

## Acknowledgements

The authors thank J. Clark Lagarias for helpful discussions and Viktoria Dubinin for assistance with development of the modified BBAG purification protocol and preparation of MBL9008307 after co-expression with pre-3 MBL9008304.

This project was financially supported by grants from the Deutsche Forschungsgemeinschaft to NFD and from the Division of Chemical Sciences, Geosciences, and Biosciences, Office of Basic Energy Sciences of the U.S. Department of Energy (DE-SC0002395) to N.C.R.

## Data Sharing and Data Availability Statement

Raw data will be made available on DataDryad.

---

**Figure S1.**
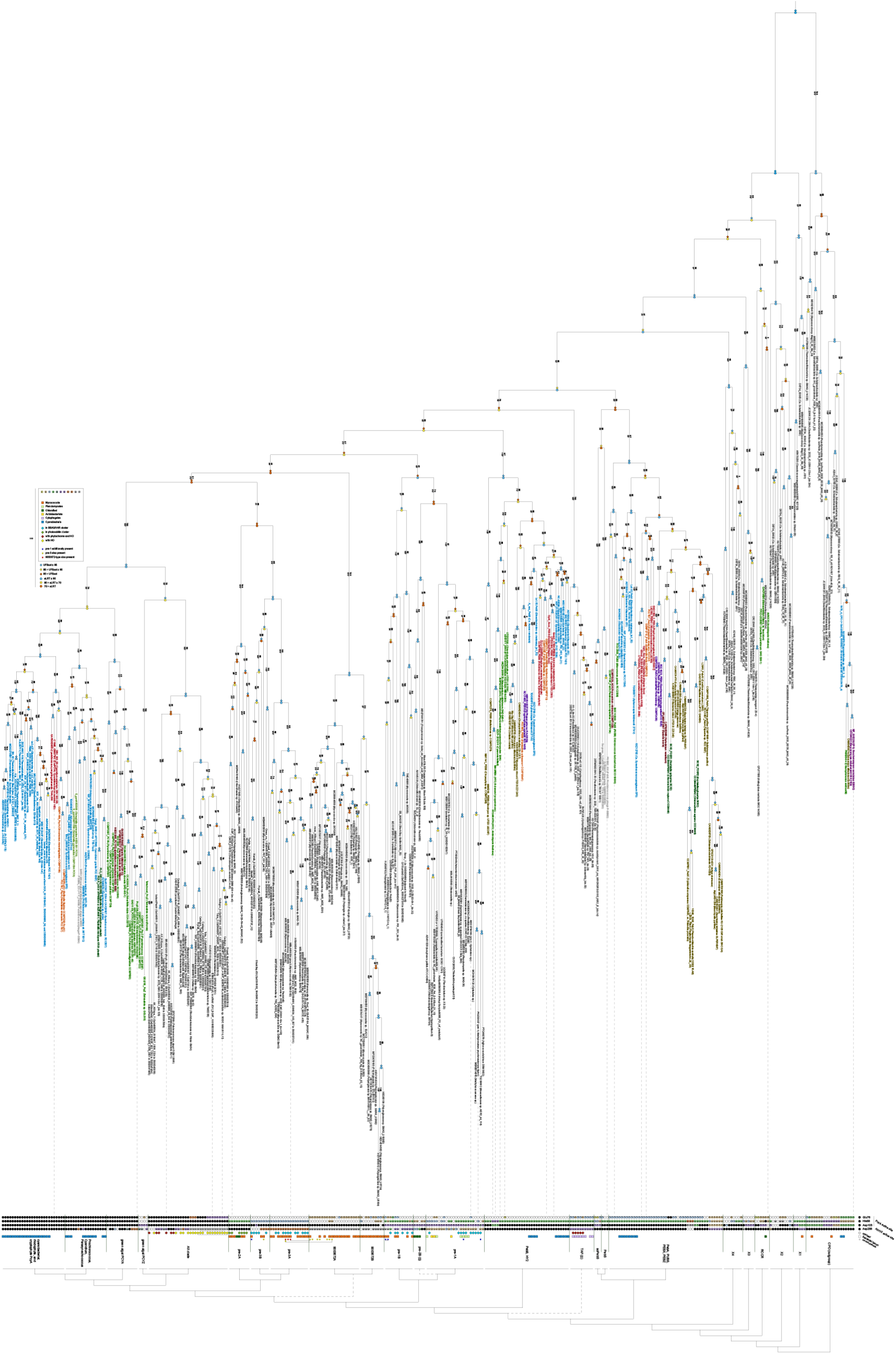
Phylogenetic analysis of FDBRs. A phylogeny was inferred for FDBRs as described in the Methods and is presented with the root between the CPO outgroup and all other sequences. Candidate active-site residues and a simple collapsed view are presented at the bottom of the figure, along with local chromosomal context. Within the collapsed tree, dashed lines indicate clades placed differently in this analysis relative to a recent analysis (Frascogna, Rockwell et al., 2026) indicating uncertain placement.

